# Differences in Protein Capture by SP3 and SP4 Demonstrate Mechanistic Insights of Proteomics Clean-up Techniques

**DOI:** 10.1101/2024.03.13.584881

**Authors:** Jessica M. Conforti, Amanda M. Ziegler, Charli S. Worth, Adhwaitha M. Nambiar, Jacob T. Bailey, Joseph H. Taube, Elyssia S. Gallagher

## Abstract

The goal of proteomics experiments is to identify proteins to observe changes in cellular processes and diseases. One challenge in proteomics is the removal of contaminants following protein extraction, which can limit protein identification. Single-pot, solid-phase-enhanced sample preparation (SP3) is a clean-up technique in which proteins are captured on carboxylate-modified particles through a proposed hydrophilic-interaction-liquid-chromatography (HILIC)-like mechanism. However, recent results have suggested that proteins are captured in SP3 due to a protein-aggregation mechanism. Thus, solvent precipitation, single-pot, solid-phase-enhanced sample preparation (SP4) is a newer clean-up technique that employs protein-aggregation to capture proteins without modified particles. SP4 has previously enriched low-solubility proteins, though differences in protein capture could affect which proteins are detected and identified. We hypothesize that the mechanisms of capture for SP3 and SP4 are distinct. Herein, we assess the proteins identified and enriched using SP3 versus SP4 for MCF7 subcellular fractions and correlate protein capture in each method to protein hydrophobicity. Our results indicate that SP3 captures more hydrophilic proteins through a combination of HILIC-like and protein-aggregation mechanisms, while SP4 captures more hydrophobic proteins through a protein-aggregation mechanism. From these results, we recommend clean-up techniques based on protein-sample hydrophobicity to yield high proteome coverage in biological samples.

## INTRODUCTION

A key goal of proteomics experiments is to identify proteins in biological samples to reveal disease biomarkers and post-translational modifications that modulate cellular activity.^1–3^ Thus, proteomics experiments require the detection and identification of proteins that have diverse properties and exist at varying concentrations in biological samples. Generally, proteomics workflows consist of protein extraction, reduction of disulfide bonds, alkylation of free cysteines, sample clean-up, protein digestion, liquid chromatography-tandem mass spectrometry (LC-MS/MS), and database searching.^4^ During protein extraction, contaminants such as buffers, salts, and chaotropic agents disrupt cell membranes to release and solubilize proteins, while maintaining physiological pH to avoid protein precipitation.^5, 6^ Though necessary for protein extraction, these contaminants suppress digestion-enzyme activity, LC resolution, and peptide ionization during electrospray, decreasing the number of identified proteins.^6, 7, 8^

Though contaminants from protein extraction must be effectively removed prior to protein digestion, sample clean-up remains a limitation in proteomics workflows.^8^ In-solution and in-gel digestions dilute contaminants rather than removing them; however, even the presence of these dilute contaminants negatively impacts the number of peptide and protein identifications.^4^ Alternative filter-based, clean-up techniques – such as filter-aided-sample preparation,^9, 10^ suspension-trapping,^11, 12^ and in-stage tip digestions^13^ – have been used to remove some contaminants,^14^ but are expensive and can result in protein loss.^15–17^

Single-pot, solid-phase-enhanced sample-preparation (SP3) is a proteomics clean-up technique that enables the identification of more peptides and proteins with higher trial-to-trial repeatability compared to the filter-based methods described above.^6, 17–20^ SP3 uses an organic, denaturing solvent to induce binding of proteins to magnetic, carboxylate-modified particles. Upon introduction of this technique, it was proposed that the proteins interacted with the particles based on a hydrophilic-interaction liquid chromatography (HILIC)-like mechanism.^21^ With the use of an external magnet, the protein-bound particles are drawn to the side of a microcentrifuge tube, allowing contaminants in the supernatant to be removed prior to protein digestion.^15^ However, the carboxylate-modified particles can (1) inadequately bind proteins, causing loss of proteins that remain in the supernatant; (2) disrupt digestion efficiency by binding to the digestion enzyme; (3) irreversibly bind proteins, preventing them from being carried into the LC-MS/MS, and (4) clog LC columns if carried into the instrument.^22–24^

The protein-capture mechanism of SP3 has recently been challenged, suggesting that instead of a HILIC-like mechanism, SP3 captures proteins primarily through a protein-aggregation mechanism.^23, 25^ Solvent precipitation, single-pot, solid-phase-enhanced sample-preparation (SP4) employs this proposed protein-aggregation mechanism and omits magnetic, carboxylate-modified particles. In SP4, an organic solvent is used to precipitate and aggregate proteins, enabling proteins to be pelleted by centrifugation. Problematic contaminants are left in the supernatant to be removed, yielding an aggregated-protein pellet for digestion. SP4 has previously enabled the identification of similar or greater numbers of peptides and proteins compared to SP3 for HEK293, Jurkat human immortalized T, and E14 murine embryonic stem-cell lysates.^22^ For HEK293 lysates, SP4 also enriched the recovery of low-solubility, transmembrane proteins compared to SP3.^22^

In proteomics workflows, identification of all proteins is ideal for profiling biological samples; thus, methods that identify proteins with differences in hydrophobicity are required. SP4 was previously shown to identify more hydrophobic proteins, yet differences in protein capture due to the carboxylate-modified particles used in SP3 could lead to the identification of different proteins. Therefore, we hypothesize that SP3 and SP4 have different mechanisms of protein capture, affecting the proteins that are identified and enriched when using each sample clean-up technique.

In this work, we aim to compare the types of proteins identified and enriched by SP3 and SP4. Additionally, we assess the ability of a sodium deoxycholate (SDC)-assisted digestion^26^ to combat the insolubility of the protein pellet generated by SP4 and increase the detection of hydrophobic proteins in both SP3 and SP4.^22^ Herein, we compare the identified proteins for samples prepared with SP3, SP4, SP3(detergent-assisted (DA)), and SP4(DA). Specifically, we use Michigan Cancer Foundation-7 (MCF7) breast cancer whole-cell lysates and subcellular fractions – including cytoplasmic, membranous, soluble-nuclear, chromatin-bound-nuclear, and cytoskeletal fractions for deep proteome profiling. Subcellular fractions are expected to contain proteins with different properties, e.g., hydrophobic, low-solubility proteins are expected in higher abundance in whole-cell lysates and membranous fractions, while hydrophilic, soluble proteins are expected in higher prevalence in cytoplasmic, soluble-nuclear, chromatin-bound-nuclear, and cytoskeletal fractions. Furthermore, subcellular fractions reduce the complexity of whole-cell lysates, often enabling the detection of more proteins, particularly those at low concentrations.^27^ Ultimately, our results show differences in the proteins identified and enriched by each clean-up technique, which we correlate to differences in protein-capture mechanisms. Finally, we recommend different sample clean-up techniques based on the expected hydrophobicity of proteins within biological samples to enable the highest number of protein identifications for effective profiling.

## EXPERIMENTAL PROCEDURES

### Materials

Ammonium bicarbonate was from Sigma Aldrich (St. Louis, MO). Emplura absolute ethanol and high-purity sodium deoxycholate (99 %) were from VWR International (Radnor, PA). Trypsin/Lys-C Mix (mass spectrometry grade) was from Promega (Madison, WI). Glu-1-Fibrinopeptide B was from Waters (Milford, MA). All other items were purchased from ThermoFisher Scientific (Waltham, MA). Nanopure water was obtained from a Purelab Flex 3 water purification system (Elga, Veolia Environment S. A., Paris, France).

### Cell culture and lysis

Epithelial estrogen receptor positive (ER+) MCF7 breast cancer cells (ATCC HTB-22, Manassas, Virginia) were cultured in Dulbecco’s Modified Eagle Medium (Corning, Corning, NY) supplemented with 10 % fetal bovine serum (Gibco, Billings, MT) and 1 % penicillin/streptomycin (Gibco, Billings, MT). Cells were regularly tested for mycoplasma contamination using PlasmoTestTM Mycoplasma Detection Kit (InvivoGen, San Diego, CA) and passaged when 80 % confluent.

Whole-cell protein was harvested using the Pierce Co-Immunoprecipitation (Co-IP) Kit (ThermoFisher Scientific, Waltham, MA) and subcellular protein fractionation was performed on 1.0 x 10^7^ cells using the Subcellular Protein Fractionation Kit for Cultured Cells (ThermoFisher Scientific, Waltham, MA). Protein concentrations were determined using bicinchoninic acid (BCA) assays^28^ and subcellular fractionation was confirmed by immunoblotting with established protein markers for each cellular fraction (Figure S1 and Table S1).

### Reduction and alkylation

Protein samples were added to low-bind, 1.5-mL microcentrifuge tubes and proteins were denatured for 5 minutes at 100 °C on a Reacti-Therm Single-Block Heating module (ThermoFisher Scientific, Waltham, MA). At room temperature, dithiothreitol (DTT) was added to the protein sample (1:10 protein-to-DTT molar ratio) and the sample was incubated at 60 °C and 1,000 rpm for 30 minutes on an Eppendorf ThermoMixer Temperature Control Device (Eppendorf, Hamburg, Germany). Iodoacetamide (IAA) was then added (1:100 protein-to-IAA molar ratio) and the sample was incubated for 30 minutes at room temperature in the dark to alkylate free cysteines. The alkylation reaction was quenched with additional DTT (5:1 DTT-to-IAA molar ratio).

### Single-pot, solid phase-enhanced sample-preparation (SP3)

Speedbead magnetic carboxylate-modified particles (GE45152105050250 and GE65152105050250, GE Healthcare, Chicago, IL) were combined in a 1:1 (v/v) ratio, washed three times with water, and resuspended in water at a final concentration of 50 mg/mL. Prepared particles were added to the protein sample in a 10:1 (w/w) particle-to-protein ratio, and chilled acetonitrile (4 °C) was added in a 4:1 (v/v) acetonitrile-to-protein ratio to induce binding to the particles (Figure S2). The protein sample was incubated on a Thermomixer at 24 °C and 1,000 rpm for 5 minutes to enhance particle-protein binding, then placed on a 1 T custom-magnetic rack for 5 minutes at room temperature. Magnetic particles with bound proteins were pulled to the side of the microcentrifuge tube and contaminants in the supernatant were removed as waste. The protein-particle mixture was washed with 80 % ethanol in water (v/v), placed on the magnetic rack for 5 minutes, and the contaminant-containing supernatant was removed as waste.^15^

### Solvent precipitation, single-pot, solid-phase-enhanced sample-preparation (SP4)

Chilled acetonitrile (4 °C) was added in a 4:1 (v/v) acetonitrile-to-protein ratio to insolubilize and aggregate proteins (Figure S2). The sample was vortexed (< 500 rpm) for 5 seconds, then centrifuged for 10 minutes at 16,000 g to pellet the proteins using a Fisherbrand accuSpin Micro 17 microcentrifuge (ThermoFisher Scientific, Waltham, MA). The contaminant-containing supernatant was removed as waste, and the protein pellet was washed with 80 % ethanol in water (v/v), centrifuged for 10 minutes at 16,000 g, and the contaminant-containing supernatant was removed as waste.^22^

### Standard digestion

Trypsin/Lys-C was added in a 1:50 (w/w) enzyme-to-protein ratio with 100 mM ammonium bicarbonate, pH 8. Proteins were digested on a ThermoMixer at 37 °C and 1,000 rpm for 18 hours. Peptide samples were then either placed on a magnetic rack (SP3) or centrifuged at 16,000 g (SP4) for 10 minutes and the peptide-containing supernatant was moved to a clean sample vial.

### Detergent-assisted digestion

Trypsin/Lys-C was added in a 1:50 (w/w) enzyme-to-protein ratio with 100 mM ammonium bicarbonate (pH 8) with 1 % SDC (using a 10 % SDC (w/v) stock solution). Proteins were digested on a Thermomixer at 37 °C and 1,000 rpm for 18 hours. Acetonitrile was added to a final solvent mixture of 20 % (v/v) to increase hydrophobic peptide recovery and trifluoroacetic acid was added to 0.5 % to precipitate SDC and quench the digestion (Figure S3).^26^ Peptide samples were centrifuged for 10 minutes at 16,000 g to pellet the detergent and any insoluble debris. For SP3(DA), samples were then placed on the magnetic rack for 5 minutes to ensure particles were not carried into the instrument. The peptide supernatant for SP3(DA) and SP4(DA) samples were then moved to clean sample vials. All four sample clean-up techniques are summarized in Figure S4.

### Liquid chromatography-tandem mass spectrometry (LC-MS/MS)

Peptides were diluted in 99 % water, 0.9 % acetonitrile, and 0.1 % formic acid to a concentration of ∼0.5 µg/µL based on Thermo Scientific Nanodrop One (Waltham, MA) measurements (Figure S5). Peptides (2 µL) were injected onto a Waters nanoACQUITY UPLC (Milford, MA) with a Waters ACQUITY UPLC M-Class Symmetry C18 Trap Column (5 µm, 180 µm x 20 mm) (Milford, MA) coupled to a Waters ACQUITY UPLC PST BEH C18 nanoACQUITY analytical column (1.7 µm, 75 µm x 100 mm) (Milford, MA). A gradient composed of solvent A (99.9 % water and 0.1 % formic acid) and solvent B (99.9 % acetonitrile and 0.1 % formic acid) was used to separate peptides prior to MS analysis. The full LC gradient is described in Table S2.

Peptides eluting from the LC column were sprayed into a Waters Synapt G2-S High-Definition Mass Spectrometer (Milford, MA) via the Waters Z-Spray NanoLockspray Ionization Source (Milford, MA) using Fossilion Sharp Singularity LOTUS nESI emitters (Fossiliontech, Madrid, Spain). Glu-1-Fibrinopeptide B was used as the reference lock mass with an exact *m/z* of 785.8421 for mass calibration and mass corrections post-acquisition. In positive-ion, resolution mode, the following tandem-MS parameters were used for data-independent analysis: mass resolution: 20,000, scan range: 50 *m/z* - 2,000 *m/z*, transfer collision energy ramp for ultra definition MS^E^ (UDMS^E^)^29^: mobility bins 0 – 20, hold 17 eV; mobility bins 20 – 110, increase collision energy from 17 eV to 45 eV; mobility bins 110 – 200, increase collision energy from 45 eV to 60 eV. Optimized MS parameters are shown in Figure S6 and Table S3.

### Database searching

Progenesis QI for proteomics (version 4.2, Nonlinear Dynamics, Waters Corporation) was used to search experimental data against the Uniprot Homo Sapien Reference Proteome (UP000005640), containing 80,027 proteins (canonical and isoform, downloaded June 2022). To determine the number of peptides and proteins identified by only one method, data from a single clean-up technique was searched. Whereas pairwise comparisons (SP3 versus SP4, SP3 versus SP3(DA), and SP4 versus SP4(DA)) were searched to determine enriched proteins by each method for volcano plot, GRAVY, and GO-Slim analyses. Data was imported, corrected for Glu-1-Fibrinopeptide B lock mass, and an ion accounting workflow was enabled to identify peptides using UDMS^E^. The following search parameters were used: peptide and fragment tolerances: automatic, enzyme specificity: cleavage of the C-terminus of lysine and arginine (/K-\P/R-\P), maximum ion charge: 20, missed cleavages: maximum 2, run alignment: automatic, peak picking sensitivity: automatic, maximum protein mass: 250 kDa, and ion matching requirements: 5 fragments/protein, 2 fragments/peptide, and 1 peptide/protein. The false discovery rate was set to < 1 %, with iadbs.exe in Progenesis Qi for proteomics creating randomized sequences “on the fly”. Modifications included fixed carbamidomethylation at cysteine, variable oxidation at methionine, and variable phosphorylation at serine, threonine, and tyrosine. Automatic processing was completed with relative quantitation using Hi-3 with protein grouping to generate raw and normalized protein abundances, max fold changes, and ANOVA values. All LC-MS/MS raw data have been deposited into the MassIVE and ProteomeXchange repositories with the following accession numbers: (MSV000094130) and (PXD049965).

### Statistics

For each biological sample (whole-cell lysate, cytoplasmic, membranous, soluble-nuclear, chromatin-bound-nuclear, and cytoskeletal fractions), three sample-preparation replicates were prepared for each clean-up technique (SP3, SP4, SP3(DA), and SP4(DA)), totaling 72 samples. Three technical instrument replicates were completed per sample, totaling 216 instrument runs. Microsoft Excel (Version 2301) was used for all tables, charts, and bar graphs unless stated otherwise. Statistical significance for pairwise comparisons (SP3 versus SP4, SP3 versus SP3(DA), and SP4 versus SP4(DA)) was determined by two-tailed F-tests and *t* tests. Bar graphs represent average values with error bars showing the standard error of the mean. Significance thresholds are represented by *p < 0.05, **p < 0.01, and ***p < 0.001. Identified proteins were manually searched in Excel by their description to determine whether they were identified in one or more sample clean-up techniques. Lucidchart (lucid.co) was used to prepare Venn diagrams. DAVID functional annotation tool^30^ (Version 2021, accessed February 2023) was used to map identified proteins to cellular components. All DAVID input parameters are described in Figures S8 and S9. Volcano plots were constructed in Excel using max-fold change and P-values (ANOVA) output from Progenesis to observe enriched proteins by each sample clean-up technique. Grand Average of Hydropathy (GRAVY) analysis was completed using GRAVY Calculator (https://www.gravy-calculator.de/index.php) for enriched proteins from volcano plots. PANTHER^31^ (Version 17.0, accessed February 2023) was used for GO-Slim cellular component analysis of enriched proteins from Progenesis. All PANTHER input parameters are described in Figure S10.

## RESULTS AND DISCUSSION

### Increases in peptide and protein identifications are observed for different sample clean-up techniques depending on the subcellular fraction

Proteomics experiments ideally enable the identification of all proteins within a biological sample to observe disease biomarkers and post-translational modifications.^1^ Thus, proteins with varying degrees of hydrophobicity must be captured from biological samples for detection and identification. Hydrophilic proteins are important because they play fundamental roles in cellular processes – such as intracellular transport and gene regulation.^32^ Proteins that are hydrophobic, such as membranous proteins, allow for ion movement across membranes, catalyze chemical reactions, and localize proteins to distinct organelles.^33–35^ Proteomics workflows typically yield the identification of hydrophilic proteins due to their high solubility and ionizability, allowing for better LC-MS/MS detection compared to hydrophobic proteins.^36–38^ Hydrophobic proteins are difficult to detect because solubilizing them requires detergents that are incompatible with proteomics workflows.^39^ Compared to SP3, SP4 has been shown to enrich the detection of low-solubility, transmembrane proteins, likely due to the protein-aggregation mechanism.^22^ Though it was hypothesized that SP3 also captures proteins through a protein-aggregation mechanism^22, 23^, we expect the presence of carboxylate-modified particles to enable interaction and capture of hydrophilic proteins with polar functional groups. Therefore, we hypothesize that protein capture by each sample clean-up technique is different and likely dependent on protein hydrophobicity.

One prior challenge of SP4 is the generation of a tightly aggregated protein pellet.^22^ Re-solubilization of the dense pellet can be difficult,^22^ resulting in poor tryptic digestion efficiency and lower protein sequence coverage during LC-MS/MS.^40^ Herein, we propose the use of SDC-assisted digestions to solubilize the protein pellet generated by SP4. SDC is a mild, ionic surfactant found in the gastrointestinal tract to solubilize lipid nutrients for digestion.^41^ SDC has previously been used to improve trypsin-digestion efficiency^42^ with removal by precipitation under acidic conditions prior to LC-MS/MS.^43, 44^ Additionally, SDC has enhanced the solubilization and detection of low-solubility proteins.^26, 43–48^ Thus, SDC-assisted digestions could combat the insolubility of the protein pellet generated by SP4, while also increasing detection of hydrophobic proteins in both SP3 and SP4.

We began by comparing the number of identified peptides and proteins for each clean-up method – SP3, SP4, SP3(DA), and SP4(DA) – for MCF7 cellular lysates and subcellular fractions. Subcellular fractionation was used to generate samples that are predicted to have a higher prevalence of either hydrophilic (cytoplasmic, soluble-nuclear, chromatin-bound-nuclear, and cytoskeletal fractions) or hydrophobic proteins (whole-cell lysates and membranous fractions). We tested the integrity of our fractionation method by confirming the presence of proteins with well-established localization patterns. We observed that GAPDH was detected exclusively in the cytoplasmic extract while E-cadherin was detected primarily in the membranous extract and minimally in the soluble-nuclear extract (Figure S1). Furthermore, SP1, a nuclear transcription factor was detected only in the soluble-nuclear extract, H3 was detected only in the chromatin-bound-nuclear extract, and β-actin was detected only in the cytoskeletal extract (Figure S1).

The number of peptides and proteins identified by each clean-up technique for the whole-cell lysate and membranous fraction were first compared because these samples are expected to contain proteins that are more hydrophobic compared to the other cellular fractions. On average, 11000 ± 1000, 17900 ± 400, 17000 ± 1000, and 15000 ± 600 peptides (Figure 1A) and 1260 ± 50, 1600 ± 40, 1470 ± 30, and 1340 ± 30 proteins (Figure 1B) were identified using SP3, SP4, SP3(DA), and SP4(DA), respectively, in the whole-cell lysate. SP4 enabled the identification of the highest number of peptides and proteins in whole-cell lysates when comparing all four clean-up techniques. SP4 also yielded the identification of the highest number of peptides and proteins in the membranous subcellular fraction, which includes proteins containing hydrophobic domains that interact with lipid bilayers.^38^ On average, 14000 ± 1500, 18100 ± 700, 14100 ± 200, and 16830 ± 90 peptides (Figure 1A) and 1480 ± 70, 1710 ± 40, 1290 ± 10, and 1441 ± 6 proteins (Figure 1B) were identified for SP3, SP4, SP3(DA), and SP4(DA), respectively, in the membranous fraction. Overall, SP4 yielded the highest number of peptide and protein identifications for the whole-cell lysate and membranous fraction compared to SP3 and methods using a detergent-assisted digestion.

**Figure 1.**
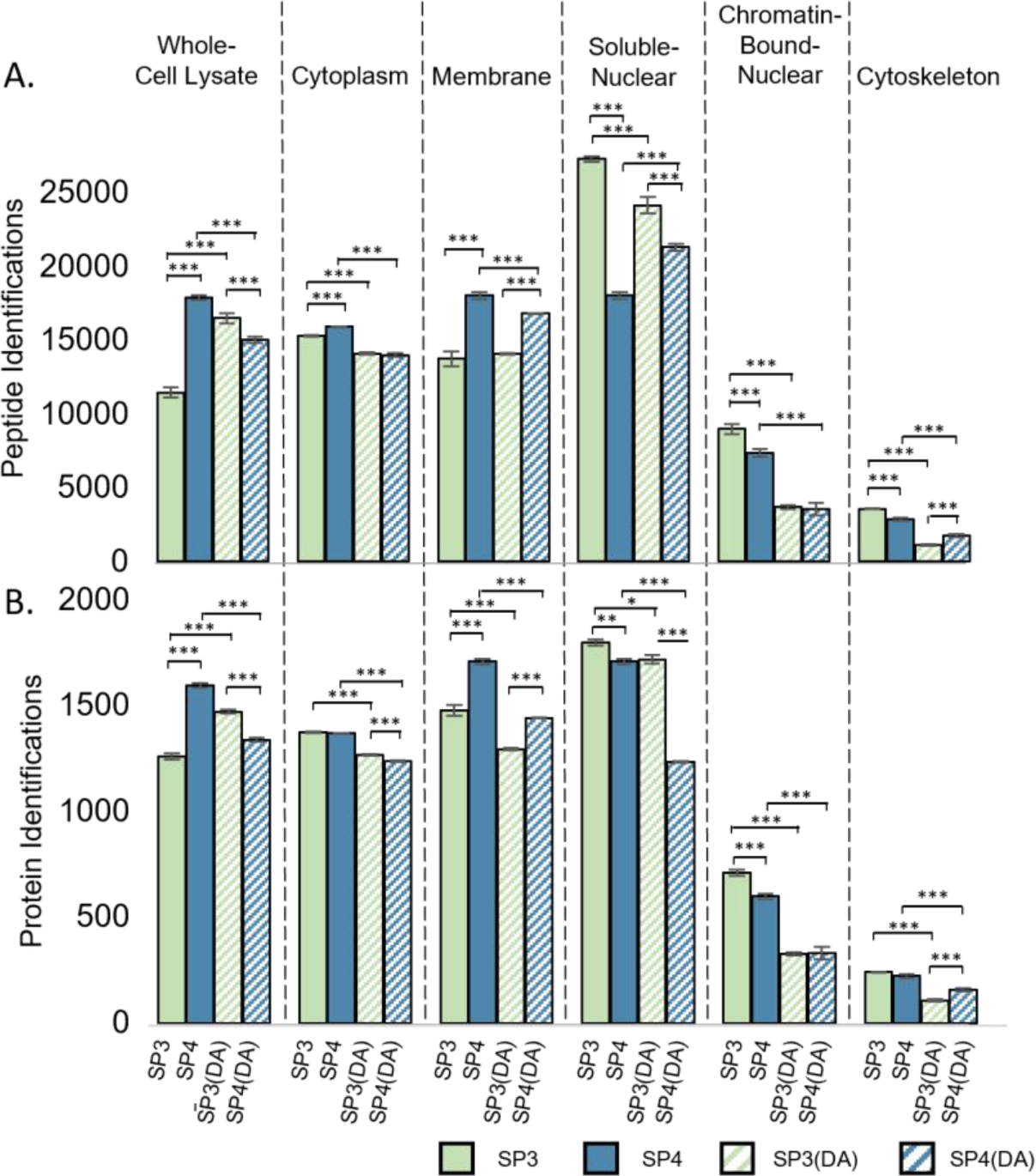
Comparisons of the average number of identified peptides (A) and proteins (B) for MCF7 breast cancer whole-cell lysates and subcellular fractions using SP3, SP4, SP3(detergent-assisted (DA)), and SP4(DA). Samples are separated by dashed lines with labels at the top of the figure. Error bars represent the standard error of the mean. Bracketed lines above bar graphs show significance of pair-wise comparisons, represented by *p < 0.05, **p < 0.01, and ***p < 0.001. Comparisons without * notation are not significantly different and are within measurement uncertainty.

We then compared the number of identified peptides and proteins for fractions that are expected to contain hydrophilic proteins, including the cytoplasmic, soluble-nuclear, chromatin-bound-nuclear, and cytoskeletal fractions.^27, 32, 49^ For the cytoplasmic fraction, on average, 15300 ± 100, 15900 ± 100, 14200 ± 300, and 14000 ± 400 peptides (Figure 1A) and 1373 ± 8, 1369 ± 6, 1270 ± 10, and 1240 ± 10 proteins (Figure 1B) were identified by SP3, SP4, SP3(DA), and SP4(DA), respectively. In the cytoplasmic fraction, SP3 and SP4 yielded the highest number of proteins, while SP4 yielded the highest number of peptides. For the soluble-nuclear fraction, on average, 27300 ± 500, 18100 ± 700, 24000 ± 2000, and 21300 ± 700 peptides (Figure 1A) and 1800 ± 40, 1710 ± 40, 1720 ± 60, and 1230 ± 10 proteins (Figure 1B) were identified by SP3, SP4, SP3(DA), and SP4(DA), respectively. SP3 yielded the identification of the highest number of peptides and proteins compared to the other methods for the soluble-nuclear fraction. For the chromatin-bound-nuclear fraction, on average, 9000 ± 1000, 7400 ± 800, 3700 ± 400, and 4000 ± 1000 peptides (Figure 1A) and 710 ± 50, 600 ± 40, 330 ± 20, and 330 ± 80 proteins (Figure 1B) were identified by SP3, SP4, SP3(DA), and SP4(DA), respectively. SP3 yielded the identification of the highest number of peptides and proteins for the chromatin-bound-nuclear fraction. For the cytoskeletal fraction, on average, 3600 ± 100, 2900 ± 300, 1100 ± 200, and 1800 ± 300 peptides (Figure 1A) and 241 ± 7, 220 ± 30, 110 ± 20, and 160 ± 20 proteins (Figure 1B) were identified by SP3, SP4, SP3(DA), and SP4(DA), respectively. SP3 yielded the identification of the highest number of peptides, while SP3 and SP4 both identified high numbers of proteins for the cytoskeletal fraction. The similarity in the number of identified proteins by SP3 and SP4 is expected when considering that both hydrophilic proteins – such as actin filaments and microtubules – and hydrophobic proteins – including intermediate filaments such as keratin, vimentin, and neurofilaments – are all present in this fraction.^27^ Trends in the numbers of identified peptides and proteins across these subcellular fractions are consistent with the increased protein recovery by SP3 compared to SP4 (Figure S5). Previously, SP3 has been shown to yield lower numbers of identified peptides and proteins in cell lysates compared to SP4.^22^ However, the results presented here demonstrate that SP3 is an effective clean-up technique for the identification of similar or higher numbers of proteins in certain subcellular fractions, specifically those that are expected to contain hydrophilic proteins at high abundance.

When assessing standard versus detergent-assisted digestions, SDC-assisted digestions did not consistently yield the highest numbers of identified peptides or proteins. When comparing SP3 to SP3(DA), the detergent-assisted digestion increased the identification of peptides and proteins in whole-cell lysates (Figures 1A and 1B). SDC-assisted digestions were used to increase both the trypsin-digestion efficiency^47^ and solubilization of hydrophobic proteins.^39^ Thus, it was expected that detergent-assisted digestions would result in higher numbers of identified peptides and proteins compared to standard digestions.^26, 50^ Though detergents can interfere with chromatographic resolution and peptide ionization,^8^ SDC was not expected to negatively affect peptide and protein identifications because it was precipitated prior to LC-MS/MS (Figure S3). Prior reports have suggested that some peptides and proteins can precipitate with SDC during acidification after digestion, decreasing the overall numbers of detected peptides and proteins.^47^ However, our results show that SDC-assisted digestions enhanced peptide and protein identifications compared to standard digestions for SP3 in whole-cell lysates, but did not yield the highest number of peptide or protein identifications when comparing all four sample clean-up techniques.

### Different sample clean-up techniques identify different types of proteins

Next, we compared the types of proteins identified by all four, or only one, clean-up technique using annotated-cellular components. All identified proteins were sorted based on whether they were identified using only one, a combination of two or three, or all four clean-up techniques (Figure 2). We used the DAVID functional annotation tool to map proteins to cellular components.^30^ The cellular components output from DAVID were manually grouped into categories: cytoplasmic, membranous, nuclear, cytoskeletal, or nonspecific components. Nonspecific cellular components included general annotations that could not be effectively grouped into the other categories, including ‘cell’, ‘cell part’, ‘intracellular organelle’, etc. Figures S8A-F and S9A-F display all cellular-component outputs from DAVID and how they were grouped into categories. Pie charts were then constructed showing the percentage of identified proteins that mapped to each cellular category (Figures 3 and S7). These analyses were performed for whole-cell lysates and each subcellular fraction.

**Figure 2.**
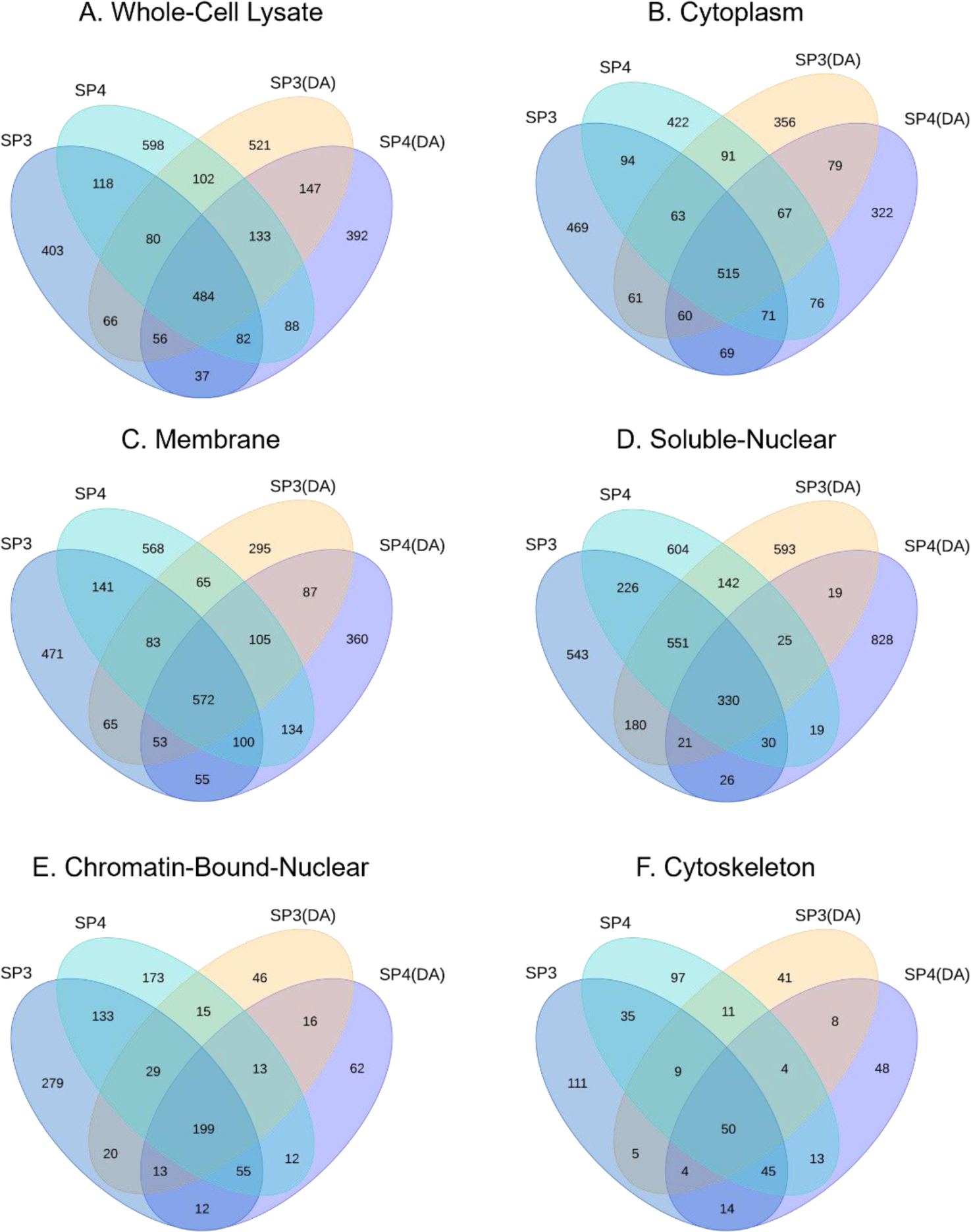
Comparison of proteins identified by each clean-up technique. Venn diagrams comparing the number of proteins identified by SP3, SP4, SP3(DA), and SP4(DA) for whole-cell lysates (A), cytoplasmic (B), membranous (C), soluble-nuclear (D), chromatin-bound-nuclear (E), and cytoskeletal (F) fractions.

**Figure 3.**
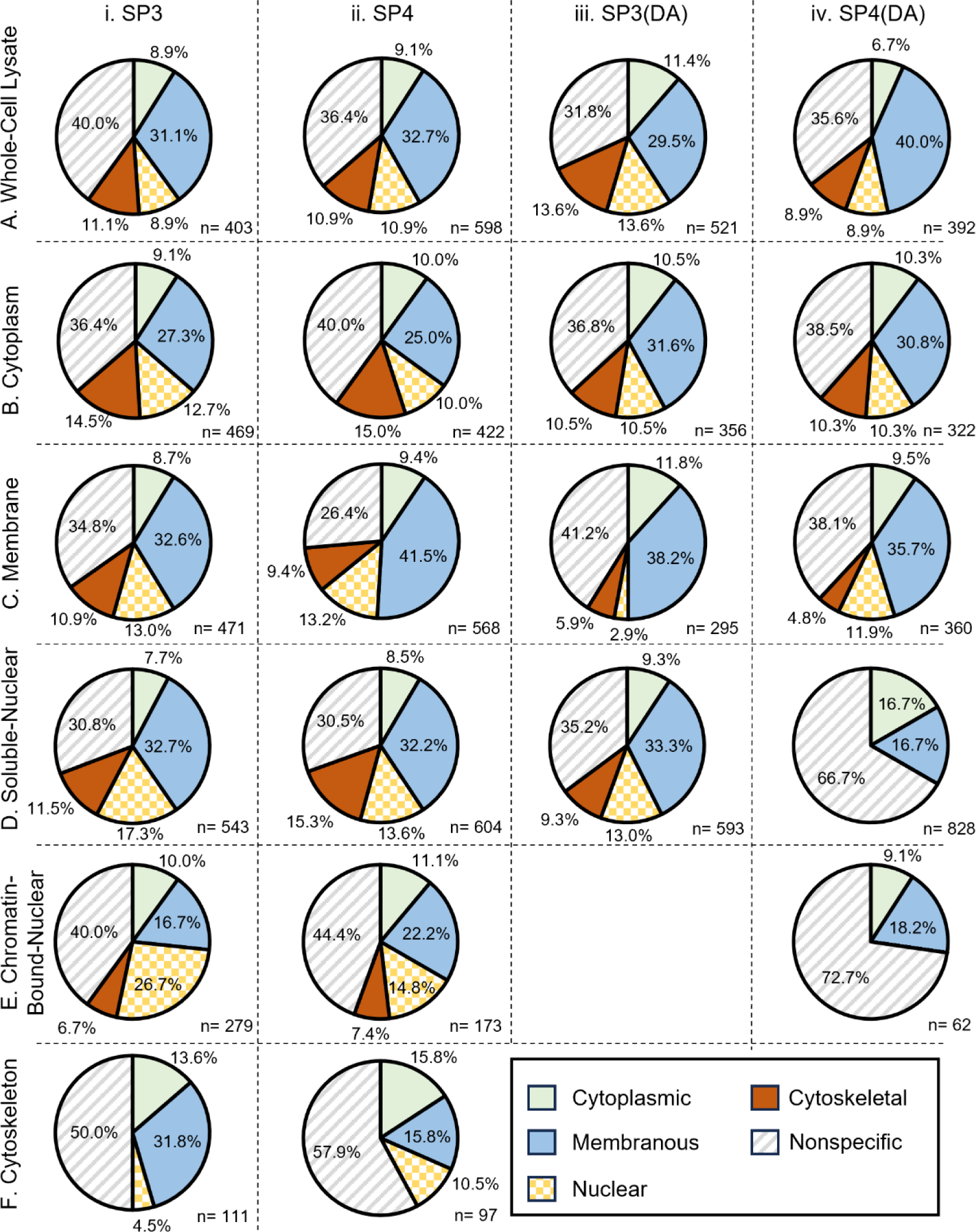
Cellular-component categories for proteins identified by only one clean-up technique (columns: SP3 (i), SP4 (ii), SP3(DA) (iii), SP4(DA) (iv)) for each sample (rows: whole-cell lysates (A), cytoplasmic (B), membranous (C), soluble-nuclear (D), chromatin-bound-nuclear (E), and cytoskeletal (F) fractions). Proteins (n) identified by only one clean-up technique were mapped to cellular components via the DAVID functional annotation tool and manually grouped into cellular-component categories: cytoplasmic (solid green), membranous (solid blue), nuclear (checkered yellow), cytoskeletal (solid red), or nonspecific (striped grey).

The proteins identified by all clean-up techniques were first assessed. There were 484, 515, 572, 330, 199, and 50 proteins identified by all four methods for whole-cell lysates, cytoplasmic, membranous, soluble-nuclear, chromatin-bound-nuclear, and cytoskeletal fractions, respectively (Figure 2). For all samples and subcellular fractions, less than 20 % of the detected proteins were identified using all four clean-up techniques, demonstrating that there are distinct differences in identified proteins based on the clean-up technique. For whole-cell lysates, the 484 proteins identified by all four clean-up techniques mapped to many different cellular-component categories, including cytoplasmic (10.1 %), membranous (40.6 %), nuclear (10.1 %), and cytoskeletal (8.7 %) (Figure S7A). This is expected because whole-cell lysates encompass proteins from all parts of the cell. Compared to whole-cell lysates, there are higher percentages of proteins mapping to membranous-cellular components in the membranous fraction (+13.6 %) (Figure S7C), nuclear-cellular components in the soluble-nuclear fraction (+10.3 %) (Figure S7D), and nuclear-cellular components in the chromatin-bound-nuclear fraction (+25.8 %) (Figure S7E). There is also a slight increase in the percentage of proteins mapping to cytoplasmic-cellular components for the cytoplasmic fraction (+0.80 %) (Figure S7B). For the cytoskeletal fraction, no cytoskeletal-grouped-cellular components were observed. This fraction is collected last in the fractionation workflow, and since proteins in different parts of the cell are known to bind to cytoskeletal proteins,^51^ it is possible that cytoskeletal proteins appear in other fractions, as seen in our results (Figures S3, S7, S8, S9). Additionally, there are proteins that mapped to cellular components in potentially unexpected fractions, *e.g.,* membranous cellular components in the cytoplasmic fraction (Figure 3 and S7). This result is in line with previous studies, which have shown that proteins can be located and distributed within multiple cellular compartments, *e.g.,* the cytoplasmic fraction can contain cytoplasmic vesicles that may map to membranous-cellular components.^32, 52–58^ Yet, these results, in combination with the Western blots presented in Figure S1, demonstrate that the subcellular fractionation was successful because for most of the subcellular fractions, increases were seen in the identified proteins associated with that cellular component.

The proteins identified using only one clean-up technique were also assessed. We first evaluated whole-cell lysates and membranous fractions, which are expected to contain hydrophobic proteins. Our results show that there were larger numbers of proteins only identified by SP4 compared to the other clean-up techniques for both of these samples (Figure 2). Proteins identified in only one clean-up technique were then mapped to cellular-component categories. Compared to all clean-up techniques, SP4(DA) yielded proteins that mapped to the highest percentage of membranous-cellular components (40 %) for whole-cell lysates (Figure 3Aiv), while SP4 yielded proteins that mapped to the highest percentage of membranous-cellular components (41.5 %) for membranous fractions (Figure 3Cii). These results indicate that for samples that are expected to contain hydrophobic proteins, there are more membranous proteins only identified by SP4 and SP4(DA) compared to SP3 and SP3(DA).

For fractions containing hydrophilic proteins (cytoplasmic, soluble-nuclear, chromatin-bound-nuclear, and cytoskeletal fractions), the proteins identified in only one clean-up technique were also assessed. There was a greater number of proteins identified by only SP3 compared to the other clean-up techniques for the cytoplasmic, chromatin-bound-nuclear, and cytoskeletal fractions (Figure 2). For the cytoplasmic fraction, SP3(DA) yielded the highest percentage of proteins mapping to cytoplasmic cellular components (10.5 %) (Figure 3Biii). For both the soluble-nuclear and chromatin-bound-nuclear fractions, SP3 yielded the highest percentage of proteins mapping to nuclear-cellular components (17.3 % for the soluble-nuclear fraction and 26.7 % for the chromatin-bound-nuclear fraction) (Figures 3Di and 3Ei). These results indicate that there are more cytoplasmic and nuclear proteins identified by only SP3 and SP3(DA) compared to SP4 and SP4(DA) for samples that are expected to contain hydrophilic proteins.

### Pairwise comparisons confirm differences in enriched proteins

Thus far, we have compared differences in the number and identity of the detected proteins using each clean-up technique. However, it is also important to compare proteins that are captured by two methods, but observed to have significantly different abundances, which we will refer to as differences in enrichment. To assess enriched proteins, pairwise comparisons (SP3 versus SP4, SP3 versus SP3(DA), and SP4 versus SP4(DA)) were used to generate volcano plots.

SP3 versus SP4 sample clean-up techniques were first compared to determine the number of proteins enriched by each method. Assessing whole-cell lysates and membranous fractions that are expected to contain more hydrophobic proteins than other cellular fractions, SP4 enriched 2.7-fold more proteins from whole-cell lysate (Figure 4Ai) and 1.8-fold more proteins from the membranous fraction compared to SP3 (Figure 4Ci). SP4 enriched more proteins in whole-cell lysates and membranous fractions compared to SP3, and this trend is consistent with the increased identifications of proteins by SP4 in these samples (Figure 1). These results are consistent with previous studies that have shown SP4 to enrich the detection of proteins in whole-cell lysates compared to SP3.^22^

**Figure 4.**
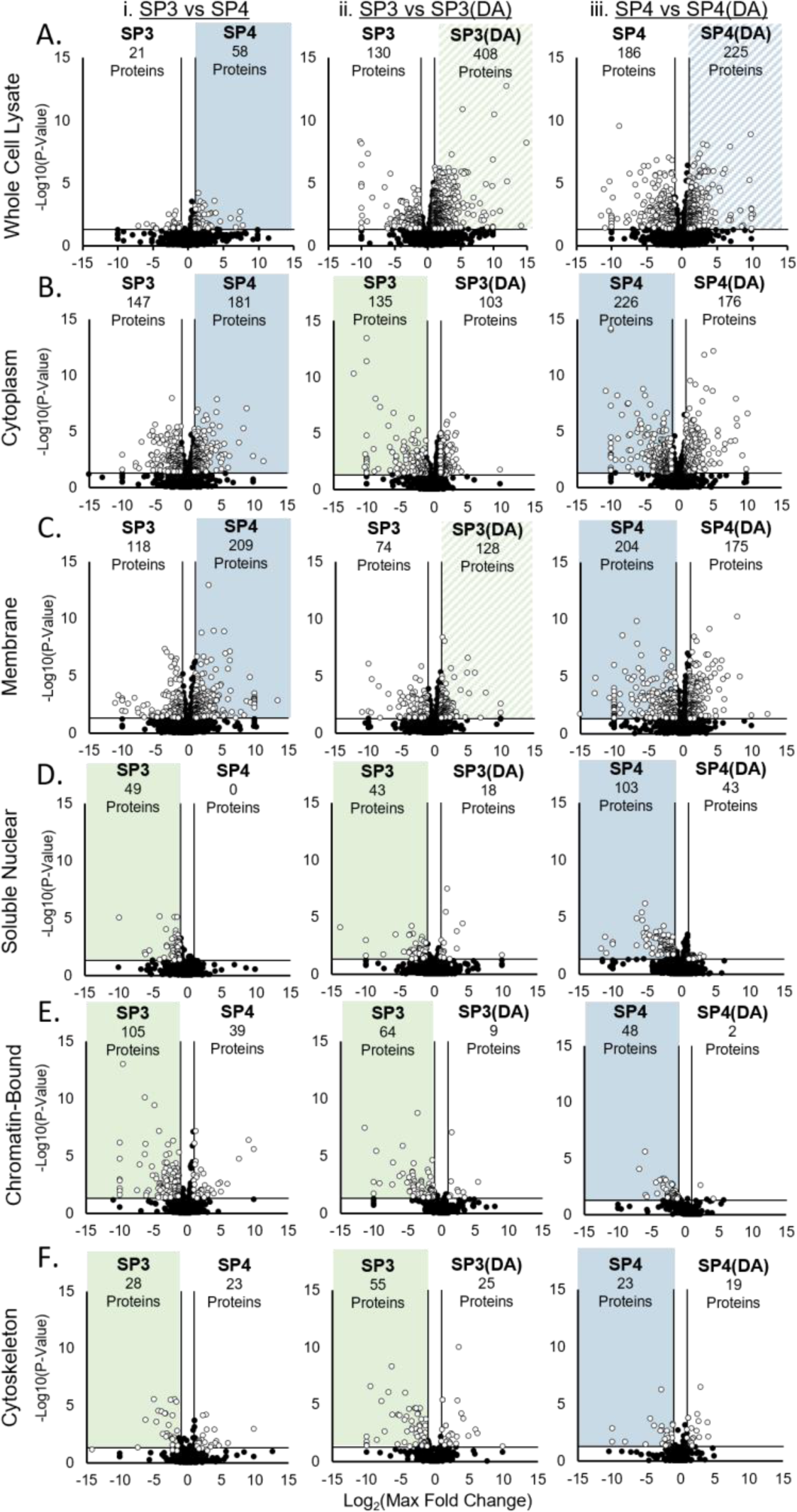
Volcano plots comparing the number of enriched proteins between SP3 versus SP4 (i), SP3 versus SP3(DA) (ii), and SP4 versus SP4(DA) (iii) in whole-cell lysate (A), cytoplasmic (B), membranous (C), soluble-nuclear (D), chromatin-bound-nuclear (E), and cytoskeletal (F) fractions. The number of enriched proteins (max fold change > 2, P-value < 0.05) for each method within the comparison is shown on the top of each panel. The highlighted panel indicates the method that yielded the highest number of enriched proteins for each pairwise comparison.

When comparing SP3 versus SP4 for subcellular fractions that are expected to contain hydrophilic proteins, SP3 enriched 2.7-fold more proteins in the chromatin-bound-nuclear fraction (Figures 4Ei) and 1.2-fold more proteins in the cytoskeletal fraction (Figure 4Fi). SP3 also enriched 49 more proteins for the soluble-nuclear fraction compared to the 0 proteins that were enriched by SP4 (Figure 4Di). For the cytoplasmic fraction, SP4 enriched 1.2-fold more proteins compared to SP3 (Figure 4Bi). However, there are many proteins enriched by both SP3 (147 proteins) and SP4 (181 proteins) (Figure 4Bi), indicating that SP3 and SP4 enrich different proteins from the cytoplasmic fraction. Thus, compared to SP4, SP3 enriches the identification of more proteins in nuclear and cytoskeletal fractions, and this trend is consistent with the increase in protein identifications within the soluble-nuclear and chromatin-bound-nuclear fractions for SP3 (Figure 1).

The number of enriched proteins was also compared for standard versus detergent-assisted digestions. For whole-cell lysates, 3.1-fold more proteins were enriched by SP3(DA) compared to SP3 (Figure 4Aii) and 1.2-fold more proteins were enriched by SP4(DA) compared to SP4 (Figure 4Aiii). In membranous fractions, 1.7-fold more proteins were enriched by SP3(DA) compared to SP3 (Figure 4Cii) and 1.2-fold more proteins were enriched by SP4 compared to SP4(DA) (Figure 4Ciii). Thus, while detergent-assisted digestions do not yield the highest number of peptide and protein identifications (Figure 1), SP3(DA) enriched more proteins compared to SP3 in whole-cell lysates and membranous fractions, while SP4(DA) enriched more proteins compared to SP4 in whole-cell lysates.

Comparing SP3 versus SP3(DA) and SP4 versus SP4(DA) in subcellular fractions that are expected to contain more hydrophilic proteins (cytoplasmic, soluble-nuclear, chromatin-bound-nuclear, and cytoskeletal fractions), standard-digestions enriched more proteins compared to detergent-assisted digestions. SP3 enriched 1.3-fold more proteins in the cytoplasmic fraction (Figure 4Bii), 2.4-fold more proteins in the soluble-nuclear fraction (Figure 4Dii), 7.1-fold more proteins in the chromatin-bound-nuclear fraction (Figure 4Eii), and 2.2-fold more proteins in the cytoskeletal fraction (Figure 4Fii) compared to SP3(DA). SP4 enriched 1.3-fold more proteins in the cytoplasmic fraction (Figure 4Biii), 2.4-fold more proteins in the soluble-nuclear fraction (Figure 4Diii), 24-fold more proteins in the chromatin-bound-nuclear fraction (Figure 4Eiii), and 1.2-fold more proteins in the cytoskeletal fraction (Figure 4Fiii) compared to SP4(DA). These results indicate that standard-digestions enrich more proteins compared to detergent-assisted-digestions for fractions that are expected to contain hydrophilic proteins.

### Proteins enriched by each clean-up technique are dependent on hydrophobicity

To this point, our data suggests that protein capture by each clean-up technique is dependent on protein hydrophobicity. To confirm that SP3 identifies and enriches proteins that are more hydrophilic while SP4 identifies and enriches proteins that are more hydrophobic, grand average of hydropathy (GRAVY) scores were calculated for the enriched proteins from volcano plots. GRAVY scores for enriched proteins were plotted as frequency distributions to compare differences in the distributions and determine whether one method enriched proteins that were more hydrophilic or hydrophobic. Proteins that are more hydrophilic or hydrophobic have GRAVY scores shifted to the more negative or positive region of the plot, respectively. Most proteins contain domains that are both hydrophilic and hydrophobic, so it is expected that GRAVY scores range between -0.2 and 0.2. Therefore, we focused on proteins with GRAVY scores outside of this range to examine differences in the hydrophobicity of proteins enriched by each method.

The GRAVY curves were first compared for SP3 versus SP4 in whole-cell lysates and membranous fractions that were expected to contain more hydrophobic proteins. For the whole-cell lysate, the GRAVY curve for SP4 has a peak apex at -0.4 and the SP3 curve has a peak apex at -0.5 (Figure 5Ai). There is also a shoulder in the more negative region of the plot for SP3 compared to SP4. In the membranous fraction, SP3 and SP4 have similar peak apexes, though, the SP3 curve has a shoulder in the more negative region of the GRAVY plot compared to SP4 (Figure 4Ci). Thus, for whole-cell lysates and membranous fractions, SP3 enriches proteins that are more hydrophilic compared to SP4.

**Figure 5.**
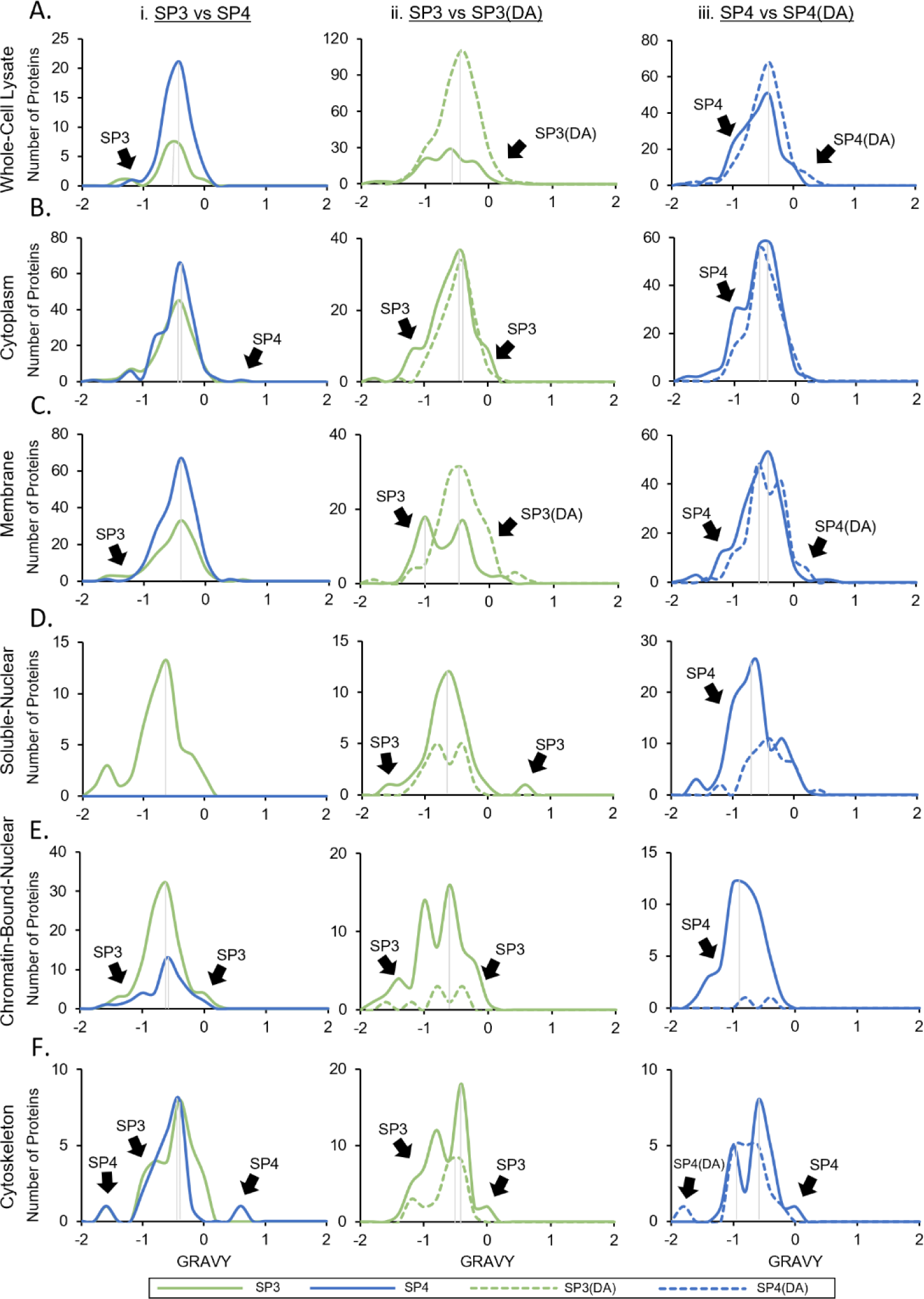
Grand average of hydropathy (GRAVY) score distributions of enriched proteins in whole-cell lysate (A), cytoplasmic (B), membranous (C), soluble-nuclear (D), chromatin-bound-nuclear (E), and cytoskeletal (F) fractions for SP3 versus SP4 (i), SP3 versus SP3(DA) (ii), and SP4 versus SP4(DA) (iii). More negative GRAVY values represent more hydrophilic proteins, while more positive values represent more hydrophobic proteins. Vertical lines indicate peak apexes to aid in comparing the distributions.

SP3 versus SP4 clean-up techniques were then compared for fractions that are expected to contain hydrophilic proteins, including the cytoplasmic, soluble-nuclear, chromatin-bound-nuclear, and cytoskeletal fractions. For the cytoplasmic fraction, the SP3 peak apex is -0.5 and the SP4 peak apex is -0.4. The GRAVY curve for SP3 is shifted to the more negative region of the plot compared to SP4, and the SP4 curve has a shoulder in the positive region of the plot with a GRAVY score of 0.5 (Figure 5Bi). For the soluble-nuclear fraction, the volcano plot showed that SP4 did not enrich any proteins compared to SP3 (Figure 4Di), so no GRAVY comparisons were made between SP3 and SP4, though, the curve for SP3 is observed in the negative region of the plot (Figure 5Di). For the chromatin-bound-nuclear fraction, the SP3 peak apex was shifted to the more negative region and there are shoulders in both the negative and positive regions of the plot (Figure 5Ei). However, the shoulder in the positive region has a GRAVY score between –0.2 to 0.2, indicating that SP3 enriched a protein containing both hydrophilic and hydrophobic domains. Our results for cellular fractions that are expected to contain hydrophilic proteins indicate that proteins enriched by SP3 are similar to or more hydrophilic compared to the proteins enriched by SP4. For the cytoskeletal fraction, the SP3 GRAVY curve extends into the positive region more than the SP4 curve, but SP4 captures one protein in the very negative region and one protein in the very positive region (Figure 5Fi). For this fraction there are lower numbers of enriched proteins compared to the other cellular fractions, and the data shows that proteins enriched by both SP3 and SP4 have attributes of hydrophilic and hydrophobic proteins. Therefore, SP3 and SP4 can be considered to enrich proteins of similar hydrophobicity for the cytoskeletal fraction. These results are consistent with the similar numbers of proteins identified by SP3 and SP4 (Figure 1B), as the cytoskeletal fraction contains both hydrophilic proteins – such as actin filaments and microtubules – and hydrophobic proteins – such as keratin, vimentin, and neurofilaments.

Previously, we assessed differences in cellular-component annotations for proteins that were identified by only one clean-up technique (Figure 3). However, analysis of the cellular-component ontologies relates enriched proteins (Figure 4) to the genes that are carrying out molecular functions and their associated localization in cells, which can provide additional insight to the biological significance of these differentially enriched proteins.^59^ Gene ontology (GO) analysis was used to determine the cellular-component ontologies of proteins enriched by each clean-up technique in pairwise comparisons. GO analysis maps proteins to cellular component ontologies by calculating the probability of observing a protein in a sample with respect to the probability of observing the protein in the entire human proteome (P-value).^31^

Enriched proteins from volcano plots for whole-cell lysates and membranous fractions were mapped to cellular-component ontologies via PANTHER to compare SP3 versus SP4, SP3 versus SP3(DA), and SP4 versus SP4(DA). GO analyses for hydrophilic fractions can be found in Figure S10. SP3 and SP4 clean-up techniques were first compared. In the whole-cell lysate, proteins enriched by SP3 mapped to cytoplasmic, membranous, nuclear, and cytoskeletal cellular-component ontologies (Figure 6Ai), while proteins enriched by SP4 mapped to membranous cellular-component ontologies (Figure 6Aii). It is expected that whole-cell lysates contain all cellular-component ontologies; thus, the fact that SP4 only enriched proteins mapping to membranous cellular-component ontologies further confirms the ability of SP4 to identify (Figures 1 and 3) and enrich (Figures 4 and 5) hydrophobic, membranous proteins. Similarly, in membranous fractions, proteins enriched by SP3 mapped to membranous and cytoskeletal cellular-component ontologies (Figure 6Bi), whereas proteins enriched by SP4 only mapped to membranous cellular-component ontologies (Figure 6Bii). These results confirm that SP3 enriches proteins mapping to cellular-component ontologies with both hydrophilic and membranous proteins, while SP4 is only observed to enrich proteins mapping to membranous cellular-component ontologies with hydrophobic proteins.

**Figure 6.**
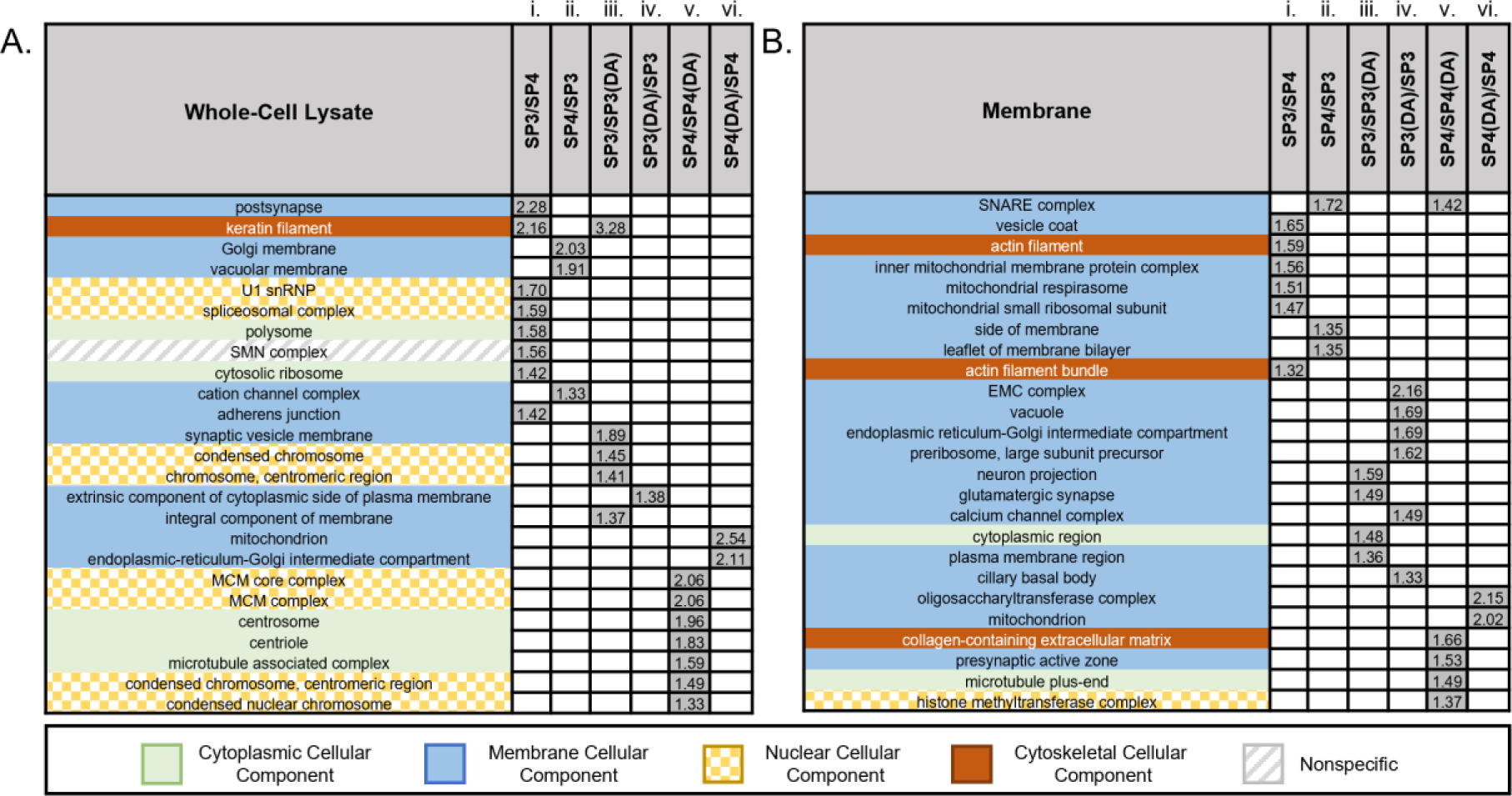
Comparison of gene ontology-Slim (GO-Slim) cellular components for enriched proteins for whole-cell lysate (A) and membranous (B) fractions. Enriched proteins were identified by volcano plots (Figure 4). Columns show comparisons of enriched cellular components for SP3 versus SP4 (SP3/SP4, i), SP4 versus SP3 (SP4/SP3, ii), SP3 versus SP3(DA) (SP3/SP3(DA), iii), SP3(DA) versus SP3 (SP3(DA)/SP3, iv), SP4 versus SP4(DA) (SP4/SP4(DA), v), and SP4(DA) versus SP4 (SP4(DA)/SP4, vi). White-colored boxes indicate that a cellular-component ontology was not significantly enriched by either method in the comparison. Grey-colored boxes represent -log_10_(P-values) for cellular components that were significantly enriched in one clean-up method versus the other via GO-Slim analysis (P-value < 0.05 in PANTHER). PANTHER input parameters are described in Figure S10 of the supporting information.

SP3 was originally proposed to capture proteins through a HILIC-like mechanism with the use of two types of hydrophilic, carboxylate-modified particles that differ in surface hydrophilicity.^15^ However, a recent study proposed that rather than HILIC-like interactions driving protein capture in SP3, protein-aggregation is primarily responsible for protein capture in both SP3 and SP4.^22^ If protein capture for SP3 was only dependent on protein-aggregation, we would expect the identified proteins in both SP3 and SP4 to yield similar protein identifications, particularly when examining subcellular fractions with reduced sample complexity. However, our results show that there are distinct differences in the proteins identified by each method (Figure 2) with less than 45 % of the identified proteins for a single fraction being identified by both SP3 and SP4. Our results show that in whole-cell lysates and membranous fractions, which are expected to contain hydrophobic proteins, SP4 enables the identification and enrichment of proteins that are similar or more hydrophobic than the proteins identified by SP3. Hydrophobic proteins contain more insoluble amino acids, which allows proteins to aggregate well with the addition of organic solvent in the protein-aggregation mechanism. These results align with prior work, which showed that SP4 enriches low-solubility proteins compared to SP3 in cell lysates.^22^ However, our results demonstrate that for fractions that are expected to contain hydrophilic proteins, SP3 enabled the identification (Figures 1, 2 and 3) and enrichment (Figures 4, 5, and 6) of proteins that are similar or more hydrophilic compared to SP4. If protein-aggregation was driving protein capture in SP3, we would expect to see similar proteins with similar levels of enrichment compared to what we observed for SP4. However, the fact that we observe more hydrophilic proteins identified with SP3 compared to SP4, suggests that HILIC-like interactions with the carboxylate-modified particles also plays a role in capturing proteins for SP3. Hydrophilic proteins have a greater affinity for polar functional groups, like carboxylates, and this affinity is based on the amount of polar and charged functional groups in the amino-acid sequence that can interact with the carboxylate group via hydrogen bonds, dipole interactions, and salt bridges.^15^ The more hydrophilic the amino-acid sequence is for the protein, the more likely it is to bind to carboxylate-modified particles. Thus, these results suggest that the protein-capture mechanisms of SP3 and SP4 are different, with protein capture for each clean-up technique being dependent on protein hydrophobicity.

GRAVY curves comparing standard and detergent-assisted digestions were also examined for whole-cell lysates and membranous fractions. In the whole-cell lysate comparison for SP3 versus SP3(DA), peak apexes were at -0.6 for SP3 and -0.4 for SP3(DA), and there is a portion of the SP3(DA) curve that is in the more positive region of the plot compared to SP3 (Figure 5Aii). For SP4 versus SP4(DA), the peak apexes for the GRAVY curves were similar, though there was a shoulder in the more positive region of the plot for SP4(DA) and a shoulder in the more negative region of the plot for SP4 (Figure 5Aiii). In comparing the membranous fraction for both SP3 versus SP3(DA) and SP4 versus SP4(DA), detergent-assisted digestions enriched proteins that appeared in the more positive regions of the plots compared to standard digestions (Figures 5Cii and 5Ciii). These results indicate that SP3(DA) and SP4(DA) increased the ability to detect more hydrophobic proteins in whole-cell lysates and membranous fractions compared to SP3 and SP4, respectively. This confirms previous studies highlighting the capabilities of detergent-assisted digestions to yield better detection of hydrophobic proteins.^39^

GRAVY curves for standard versus detergent-assisted digestions were then compared for fractions that contain hydrophilic proteins, such as cytoplasmic, soluble-nuclear, chromatin-bound-nuclear, and cytoskeletal fractions. For these fractions, GRAVY curves for standard-digestion methods had shoulders that were shifted to more negative regions of the plots compared to the curves for detergent-assisted digestions (Figures 5Bii, 5Biii, 5Dii, 5Diii, 5Eii, and 5Eiii). In the comparison of SP4 versus SP4(DA) for the cytoskeletal fraction, the GRAVY curves were similar, with SP4 enriching one protein in the positive region, and SP4(DA) enriching one protein in the negative region of the GRAVY plot (Figure 5Fiii). Thus, SP4 and SP4(DA) enrich proteins with similar hydrophobicity in the cytoskeletal fraction. Together, standard digestions appear to increase the detection of proteins that are similar or more hydrophilic compared to detergent-assisted digestions, and detergent-assisted digestions appear to increase the detection of more hydrophobic proteins compared to standard digestions.

Proteins enriched by standard digestions and detergent-assisted digestions were also assessed using GO-Slim for pairwise comparisons of SP3 versus SP3(DA) and SP4 versus SP4(DA). Enriched proteins by standard-digestions in whole-cell lysates and membranous fractions map to cytoplasmic, membranous, nuclear, and cytoskeletal cellular-component ontologies (Figures 6Aiii, 6Av, 6Biii, and 6Bv). Whereas proteins enriched by detergent-assisted digestions only map to membranous cellular-component ontologies (Figure 6Aiv, 6Avi, 6Biv, and 6Bvi). These results further confirm that detergent-assisted digestions increase the ability of SP3 and SP4 methods to detect membranous proteins in whole-cell lysates and membranous fractions compared to standard-digestions.

## CONCLUSION

Proteomics experiments enable the identification of many peptides and proteins in biologically relevant samples to determine important protein biomarkers in disease. Herein, we compare SP3, SP4, SP3(DA), and SP4(DA) and show that protein capture by each clean-up technique is dependent on protein hydrophobicity. Our results demonstrate that SP3 captures proteins through a combination of HILIC-like interactions and a protein-aggregation mechanism, while SP4 appears to only capture proteins through a protein-aggregation mechanism. Further, detergent-assisted digestions improve the identification of hydrophobic, membranous proteins in whole-cell lysates and membranous fractions.

Though an ideal proteomics method would successfully capture all proteins from a biological sample, we show that SP3 and SP4 capture different proteins. Thus, we recommend using SP3 with standard-trypsin digestions when attempting to identify the highest number of proteins in samples that are expected to contain primarily hydrophilic proteins. Alternatively, we recommend using SP4 with standard-trypsin digestions when attempting to identify the highest number of proteins in samples that are expected to contain hydrophobic proteins. Lastly, we recommend using detergent-assisted digestions when attempting to improve the detection of membranous proteins with many hydrophobic domains. Our results simplify the decision-making process for proteomics users to choose a method that identifies and enriches the highest number of proteins for different types of biological samples. By using a recommended clean-up technique, proteomics experiments may be able to achieve higher proteome coverage of complex samples, which would enable the analysis of proteins in their vital roles that modulate cellular activity and lead to disease.

## ASSOCIATED CONTENT

### Supporting Information

Western blots; Denaturing solvent analysis; SDC pelleting experiments with TFA; Summarized clean-up methods; Protein recovery by Nanodrop; Driftscope mobility; Annotated-cellular components of proteins identified in all clean-up techniques; Annotated-cellular components of proteins identified in only one clean-up technique; Gene-ontology-Slim analyses; Protein concentrations via BCA assays; Full LC gradient; Optimized MS Parameters. (PDF)

### Author Information

#### Author Contributions

J.M.C., J.H.T., and E.S.G were responsible for experimental planning. C.S.W cultured and fractionated MCF7 cells. A.M.N. and J.T.B. performed BCA assays and western blots. J.M.C. was responsible for all proteomics sample-preparation, LC-IMS-MS/MS runs, database searching, and data analysis. A.M.Z. assisted J.M.C. with sample-preparation and data analysis. J.M.C. and E.S.G wrote the manuscript, and all others approved the final version.

## ACKNOWLEDGEMENT

The authors wish to acknowledge the Baylor University Mass Spectrometry Center and Molecular Biosciences Center for resources and instrumentation. We also thank Dr. Luke T. Richardson for helpful discussions.

## ABBREVIATIONS

ANOVA: analysis of variance
BCA: bicinchoninic assay
DAVID: database for visualization and integrated discovery
DA: detergent-assisted
DTT: dithiothreitol
GAPDH: glyceraldehyde-3-phosphate dehydrogenase
GO: gene ontology
GRAVY: grand average of hydropathy
H3: histone 3
HILIC: hydrophilic interaction liquid chromatography
IAA: iodoacetamide
LC-MS/MS: liquid chromatography-tandem mass spectrometry
MCF7: Michigan cancer foundation-7
PANTHER: protein analysis through evolutionary relationships
SP3: single-pot solid phase enhanced sample preparation
SDC: sodium deoxycholate
SP4: solvent precipitation single pot solid phase enhanced sample preparation
SP1: specificity protein 1
UDMS^E^: ultra definition MS^E^
UPLC: ultra-performance liquid chromatography.

## Table of Contents Graphic

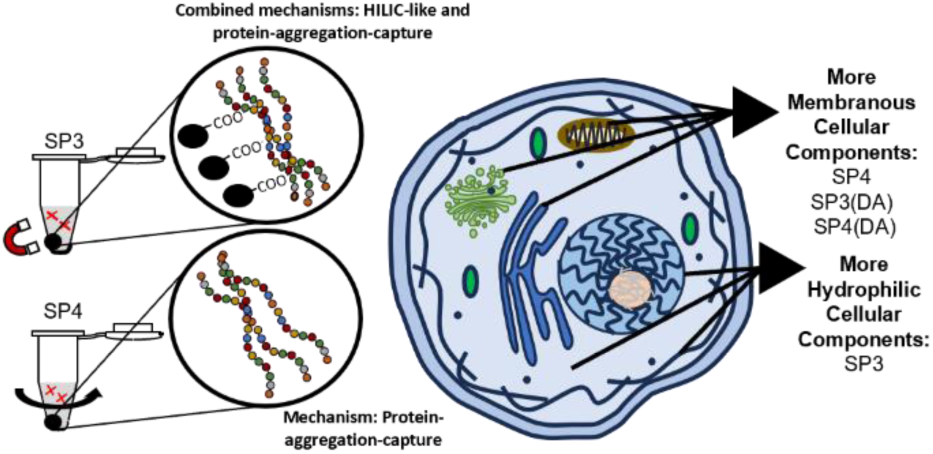

## Supporting Information

**Figure S1.**
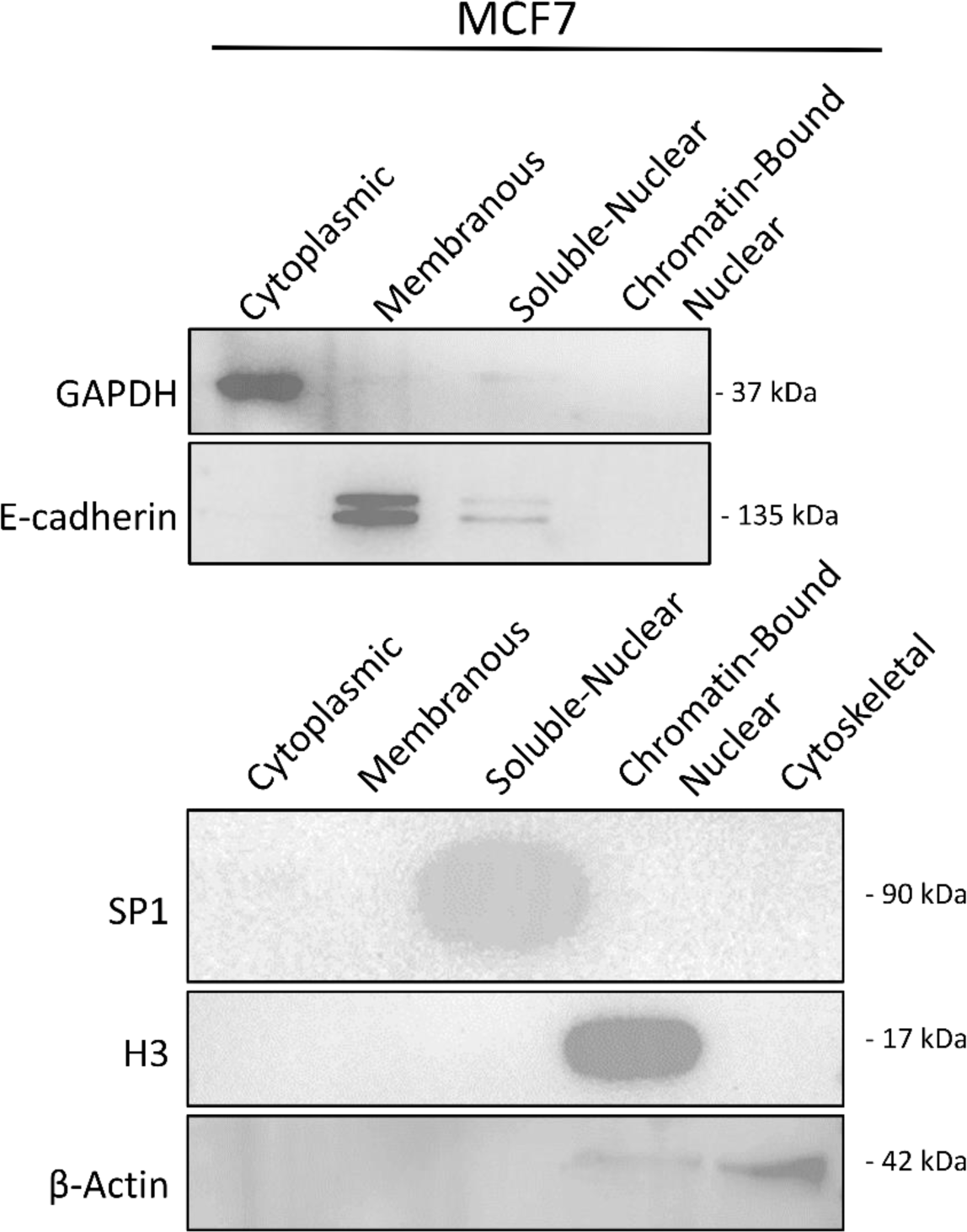
Western blot analysis confirms subcellular fractionation. Primary antibodies for proteins that exist primarily within a singular subcellular fraction (GAPDH: Cytoplasm, E-cadherin: Membrane, Specificity protein 1 (SP1): Soluble-Nuclear, Histone (H3): Chromatin-Bound Nuclear, β-actin: Cytoskeletal) were used to confirm the integrity of subcellular fractionation.

### Western Blot Methods

Proteins were separated by an SDS/PAGE gel and transferred onto a 0.45 µm PVDF membrane (ThermoFisher Scientific, Walthum, MA). Membranes were probed with primary antibodies overnight at 4 °C followed by secondary antibodies for 1 hour at room temperature. Chemiluminescence was detected with ECL Prime (Cytiva, Marlborough, MA). Blot images were captured using the ChemiDoc Imaging System (BioRad, Hercules, CA). The following antibodies were used: anti-GAPDH (Catalog No. 51332, Cell Signaling Technology, Danvers, MA), anti-E-cadherin (Catalog No. 610404, Becton Dickinson, Franklin Lakes, NJ), anti-SP1 (Catalog No. 9389S, Cell Signaling Technology, Danvers, MA), anti-H3 (Catalog No. 9715, Cell Signaling Technology, Danvers, MA), anti-β-Actin (Catalog No. 612657, Becton Dickinson, Franklin Lakes, NJ), Horseradish peroxidase (HRP) goat anti-mouse (Part No. 926-80010, LI-COR Biosciences, Lincoln, NE), and HRP goat anti-rabbit (Catalog No. 7074-S, Cell Signaling Technology, Danvers, MA).

**Figure S2.**
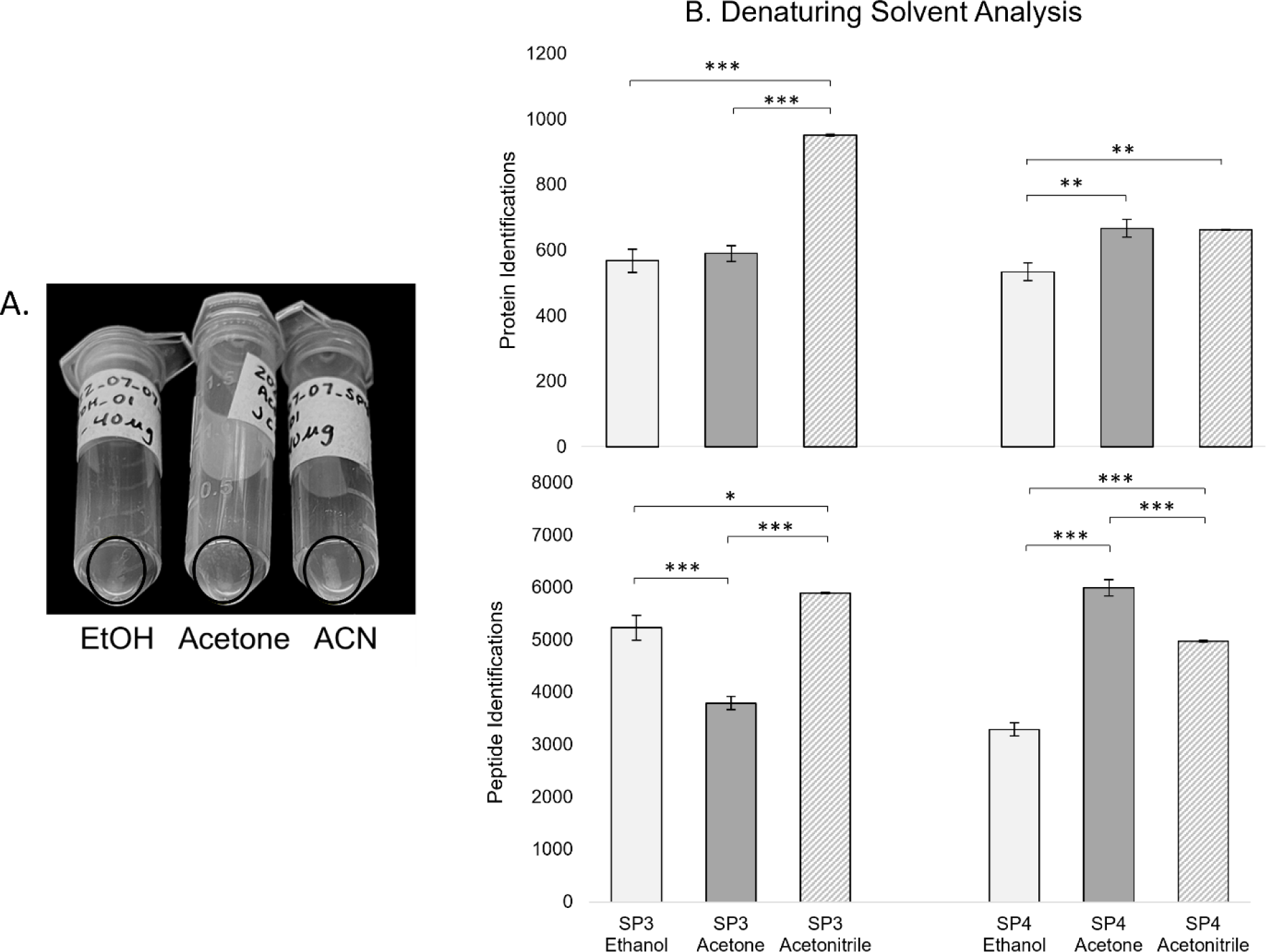
Acetonitrile is an acceptable denaturing solvent. Ethanol, acetone, and acetonitrile were assessed to determine the denaturing solvent that is most effective for aggregating proteins in SP4 (A) and yields the highest number of peptide and protein identifications for SP3 and SP4 (B). Error bars in bar graphs display the standard error of the mean. Bracketed lines above bar graphs show the significance of pair-wise comparisons (*e.g.,* ethanol versus acetone, ethanol versus acetonitrile, and acetone versus acetonitrile), represented by *p < 0.05, **p < 0.01, and ***p < 0.001. Comparisons without * notation are not significantly different and are within measurement uncertainty.

### Figure S2 Results and Discussion

Acetonitrile (ACN) displayed the largest and most opaque pellet (A), yielded the highest number of identified peptides and proteins for SP3, is one of the methods that identifies the highest number of proteins for SP4, and has the lowest variability in number of identified proteins and peptides for both SP3 and SP4 (B). Acetonitrile and acetone have previously been shown to yield similar peptide and protein identifications.^1^ But, other studies of SP3 and SP4 used acetonitrile;^1^ thus, acetonitrile was chosen as the denaturing solvent for this work so we could draw direct comparisons.

**Figure S3.**
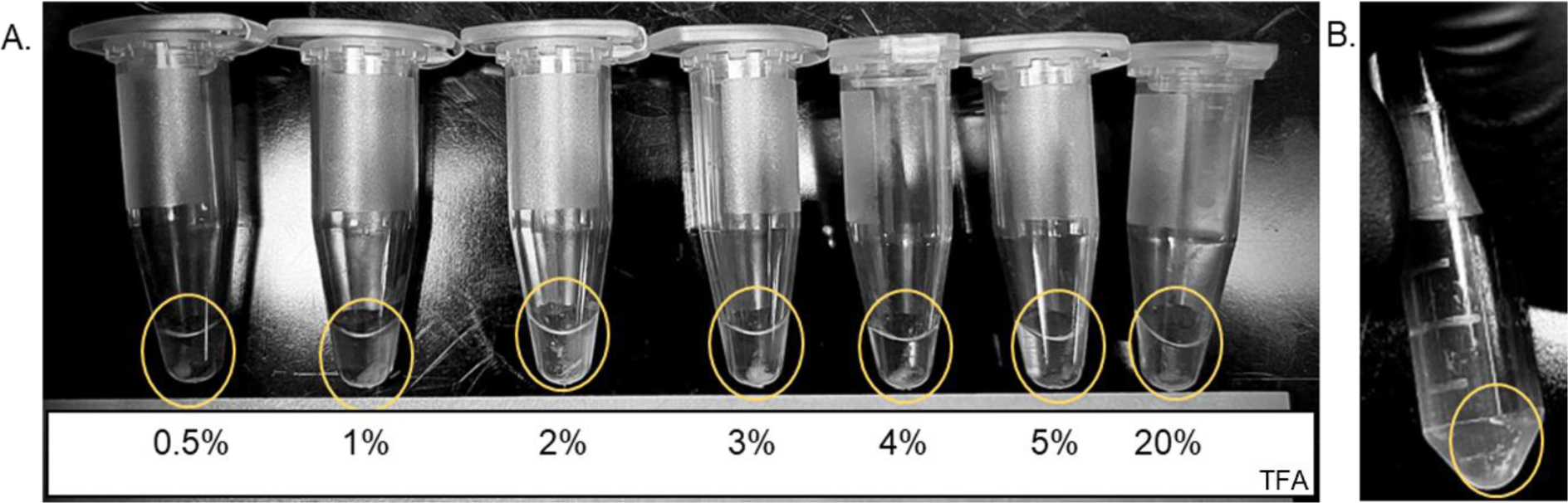
Sodium deoxycholate (SDC) is effectively pelleted with 0.5% trifluoroacetic acid (TFA). There were no observable differences in the size of the SDC pellet as the percentage of TFA was increased. Therefore, 0.5% TFA was chosen for all experiments.

### Sodium-Deoxycholate Pelleting Methods

TFA has previously been used to pellet SDC.^2^ Seven concentrations of TFA (0.5%, 1%, 2%, 3%, 4%, 5%, and 20%) were used to pellet SDC from 100 mM ammonium bicarbonate with 1% SDC, pH 8. The mixtures were centrifuged for 10 minutes at 16,000 g, and each TFA concentration was able to pellet SDC.

**Figure S4.**
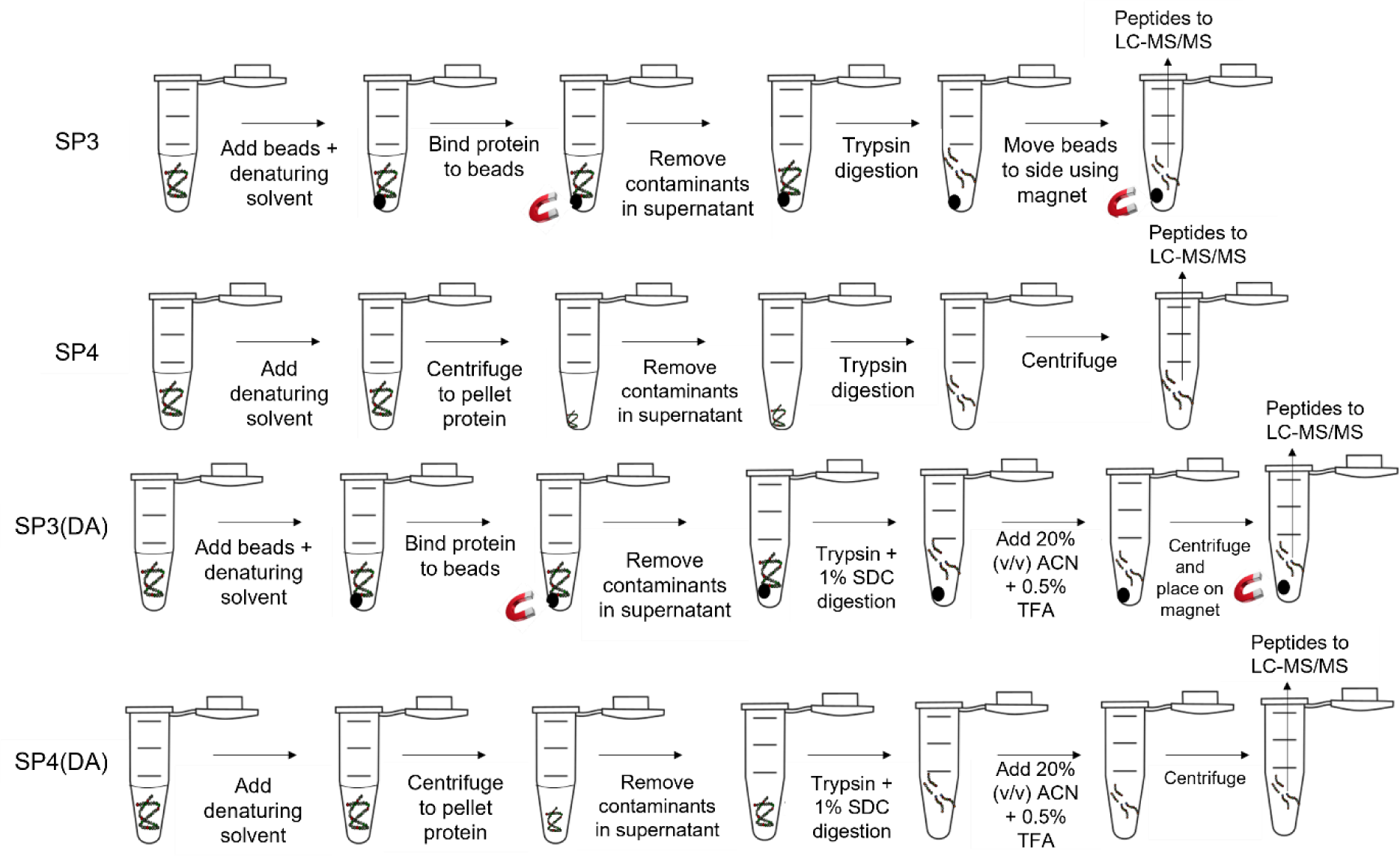
Summary of sample clean-up methods.

**Figure S5:**
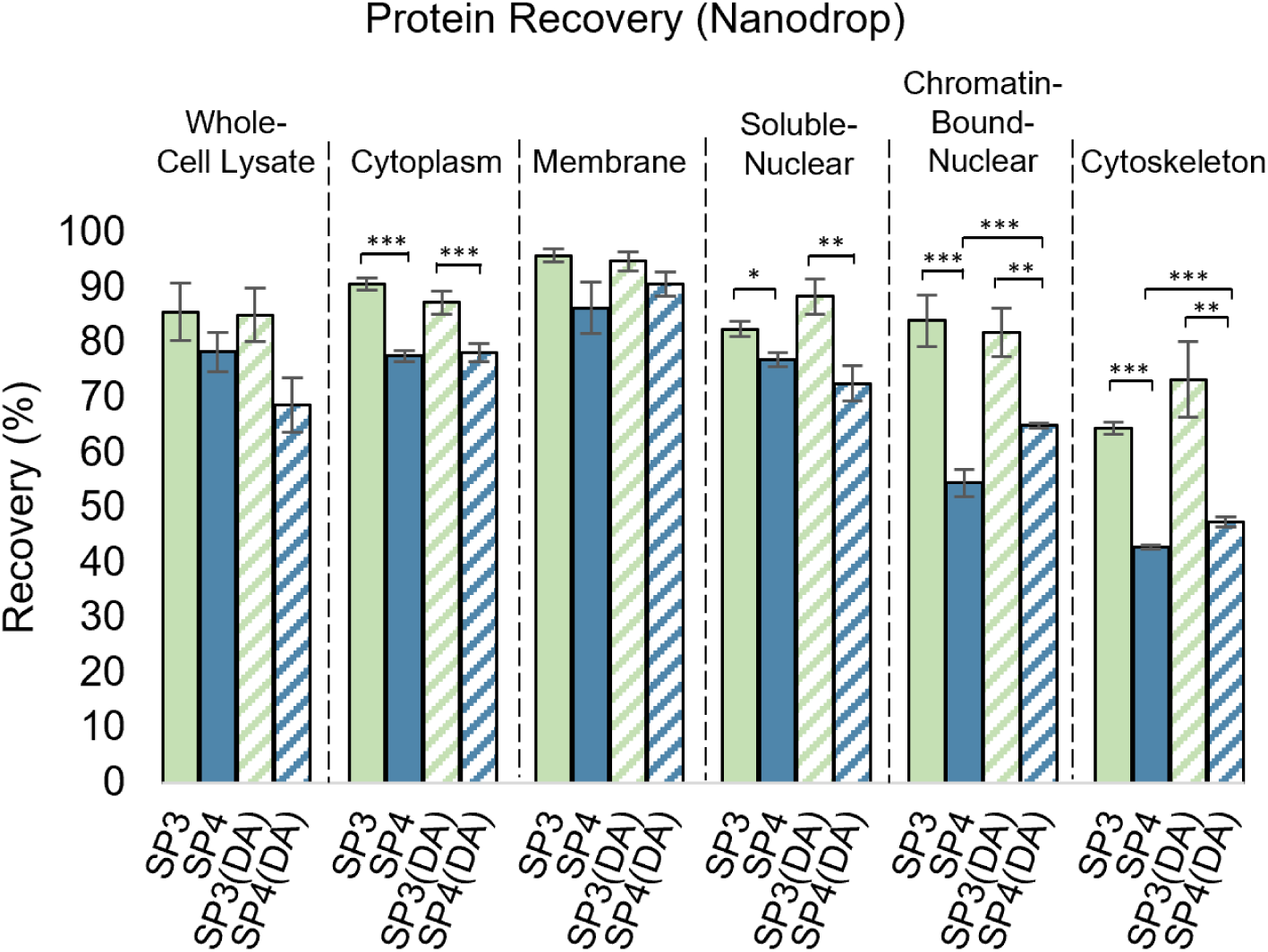
Protein recovery differs among subcellular fractions. In fractions that are expected to contain more hydrophilic proteins (*e.g.,* cytoplasmic, soluble nuclear, chromatin-bound-nuclear, and cytoskeletal fractions), SP3 yielded higher protein recovery compared to SP4. In fractions that are expected to contain more hydrophobic proteins (*e.g.,* whole-cell lysate and membrane fractions), protein recoveries for all pairwise comparisons (SP3 vs SP4, SP3 vs SP3(DA), SP4 vs SP4(DA), and SP3(DA) vs SP4(DA)) were similar. These results suggest differences in protein recovery based on sample clean-up technique.

### Protein Recovery Methods

Peptide concentrations for each clean-up technique were measured in triplicate on a Nanodrop One (ThermoFisher Scientific, Waltham, MA) prior to LC-MS/MS analysis. Protein recoveries (%) were calculated by computing the protein recovered in micrograms (Eq. S1) using the total volume (vol) (Eq. S2). Volumes of ACN and TFA were only used for detergent-assisted digestions.

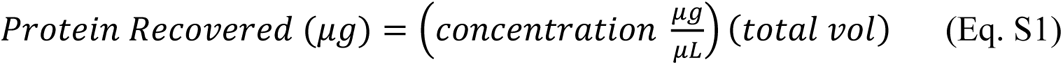

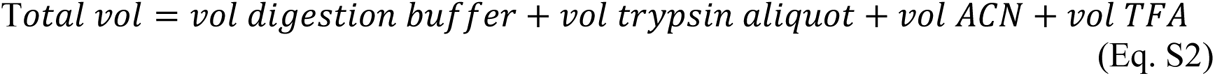

The percent protein recovery was then determined using the protein recovered (Eq. S1) and the starting protein amount, measured by a BCA assay (Eq. S3).

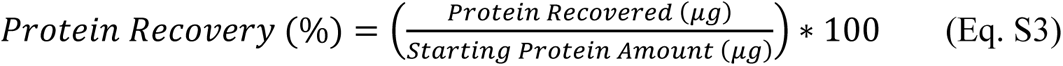

**Figure S6.**
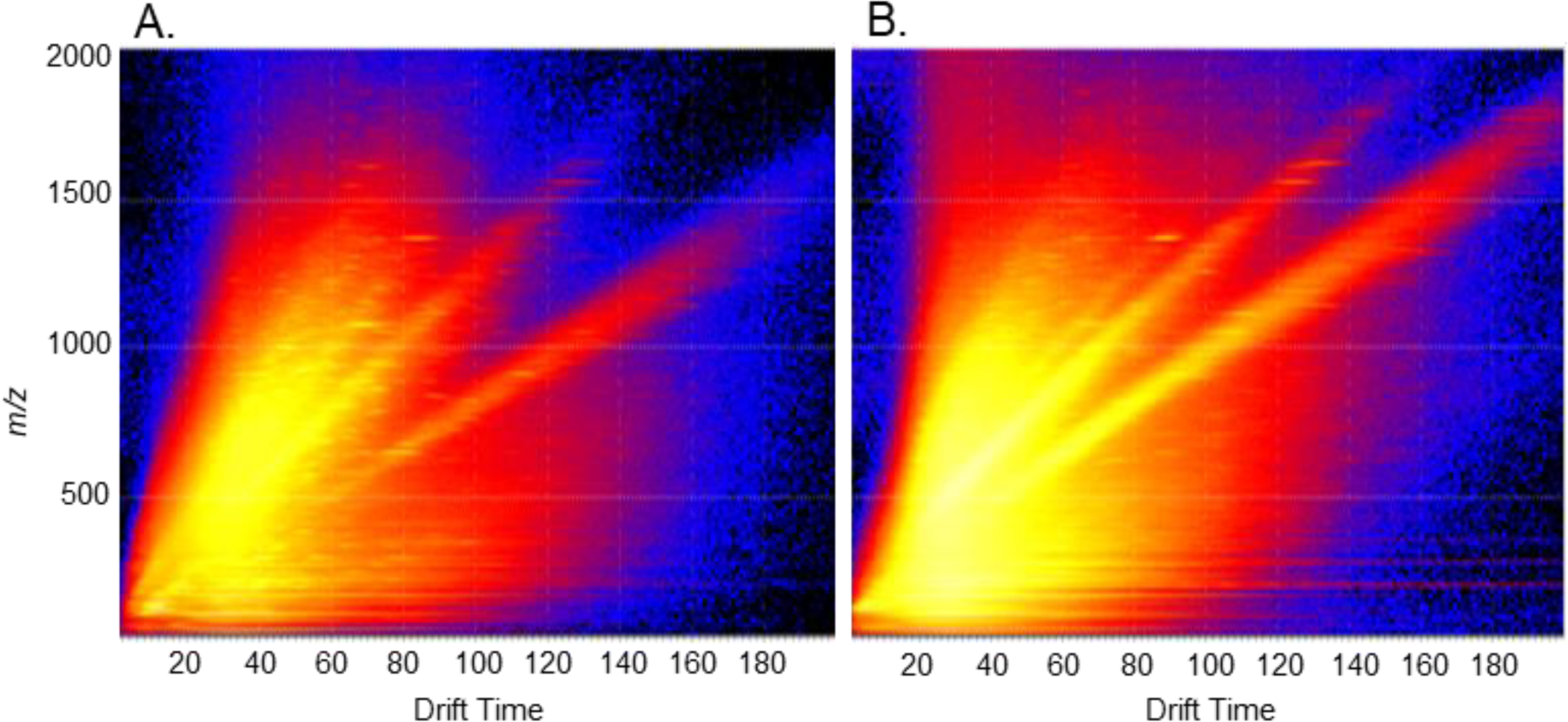
Driftscope Mobility Separation. Runs examining whole-cell lysate were input into Driftscope (Version 2.9) using Function 2 (mobility function) to visualize the separation of precursor ions in the mobility cell. Tune-page settings from previous literature^3^ (A) and after optimization (B) were compared to determine the ideal parameters for ion-mobility separations. Heat maps display the intensity of precursor ions during the mobility separation, with warm colors (yellow and red) indicating greater signal intensity. Optimized tune-page parameters (B) increase the range of drift times for the precursor ions in the mobility cell, shown by the heat map intensities extending into higher *m/z* and drift-time ranges. A full list of MS parameters can be found in Table S3.

**Figure S7.**
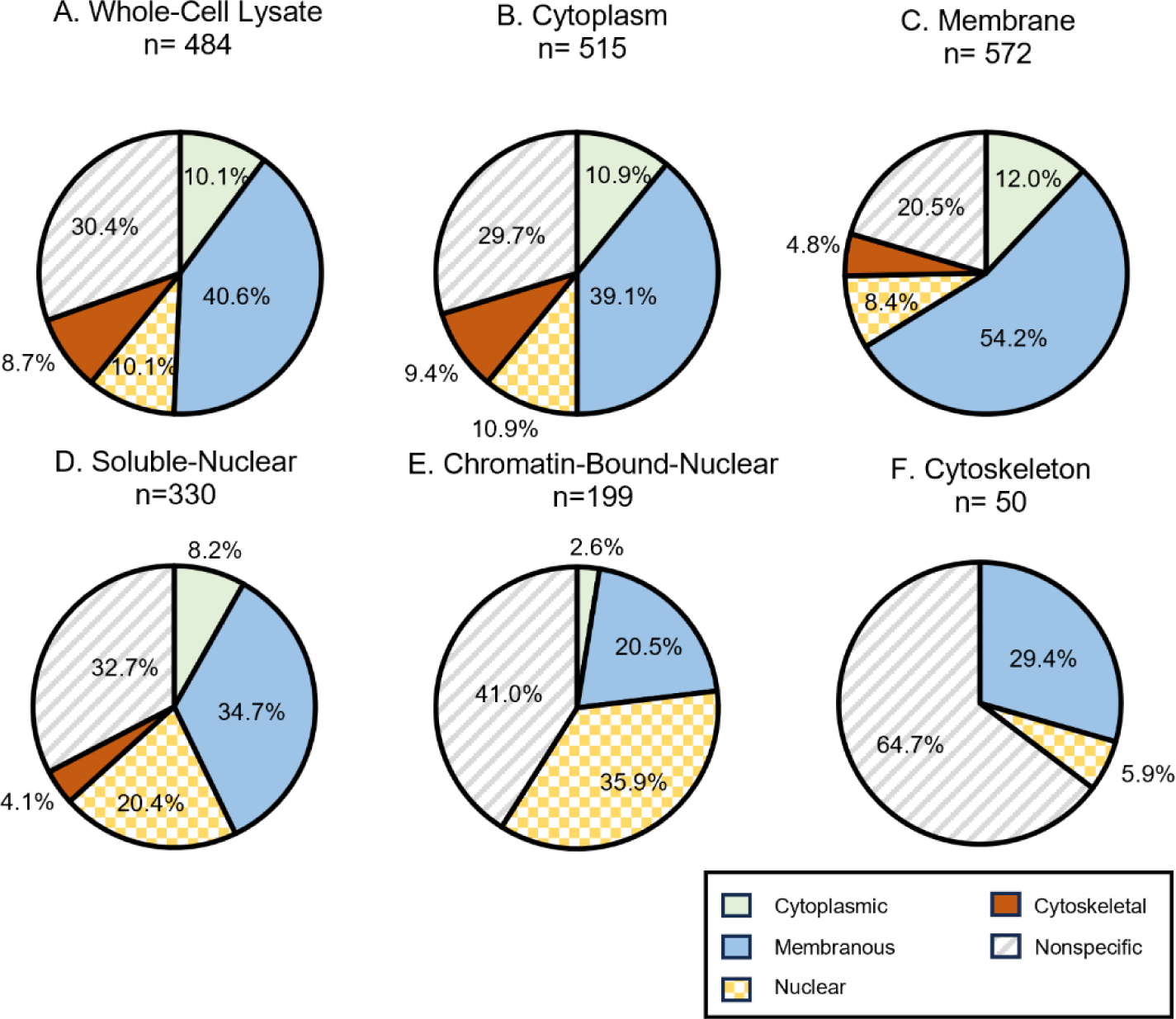
Cellular-component categories for proteins identified by all four clean-up techniques for whole-cell lysates (A), cytoplasmic (B), membranous (C), soluble-nuclear (D), chromatin-bound-nuclear (E), and cytoskeletal (F) fractions. Proteins (n) identified by all four clean-up techniques were mapped to cellular-component annotations via the DAVID functional annotation tool^4^ and manually grouped into categories: cytoplasmic (solid green), membranous (solid blue), nuclear (checkered yellow), cytoskeletal (solid red), or nonspecific (striped grey). Category percentages were calculated by dividing the number of mapped-cellular components in a category by the total number of mapped-cellular components.

**Figures S8A-F.**
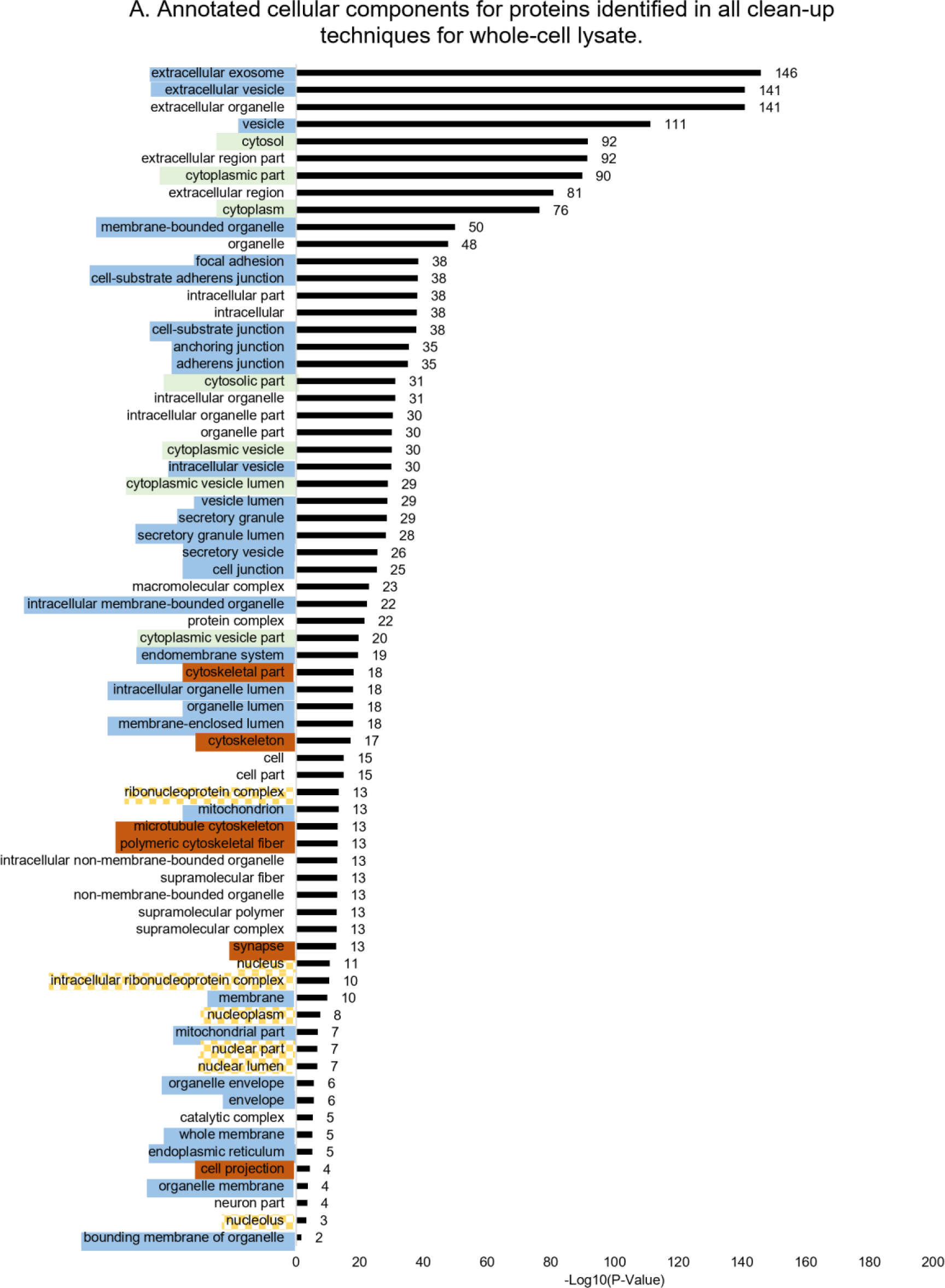
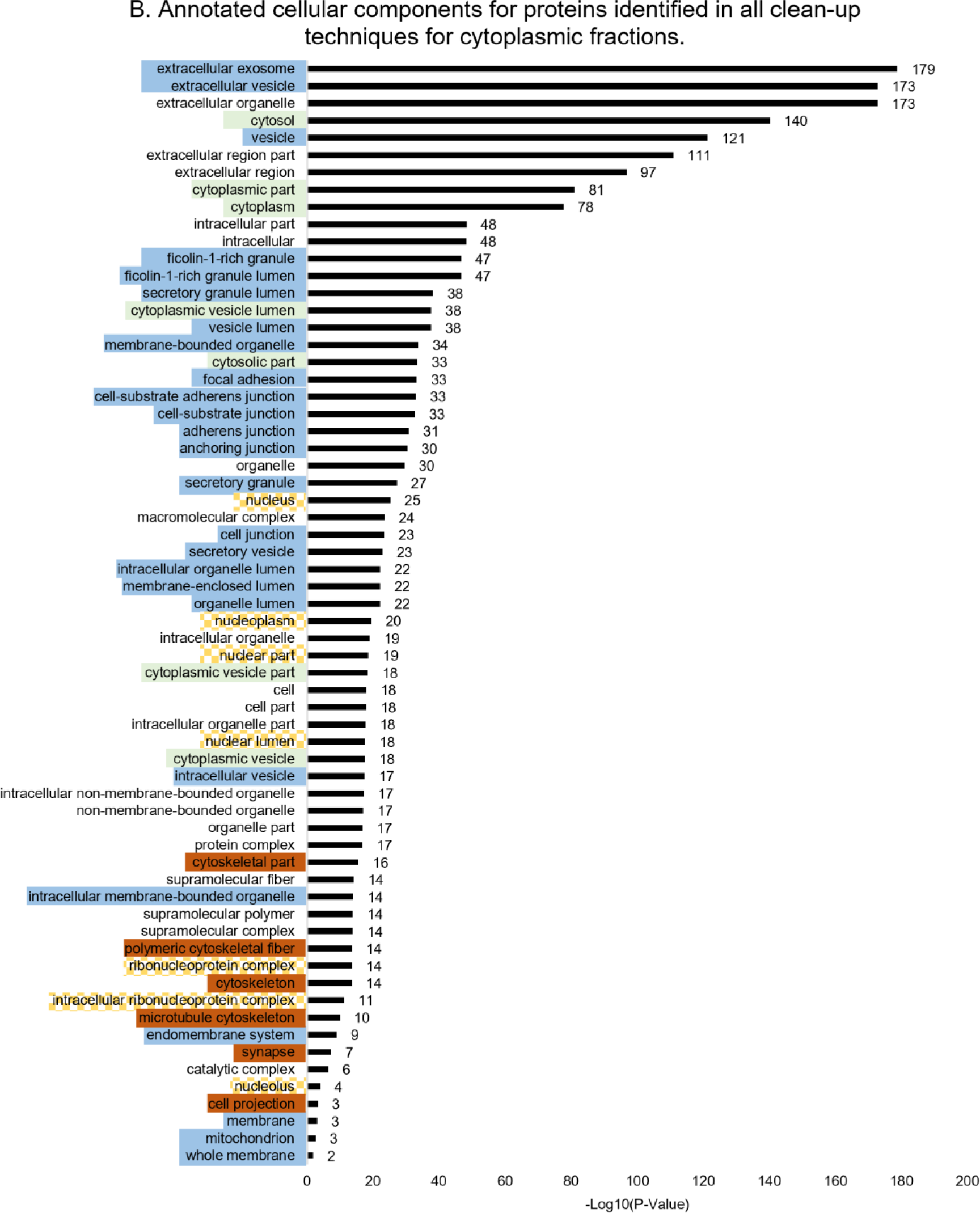
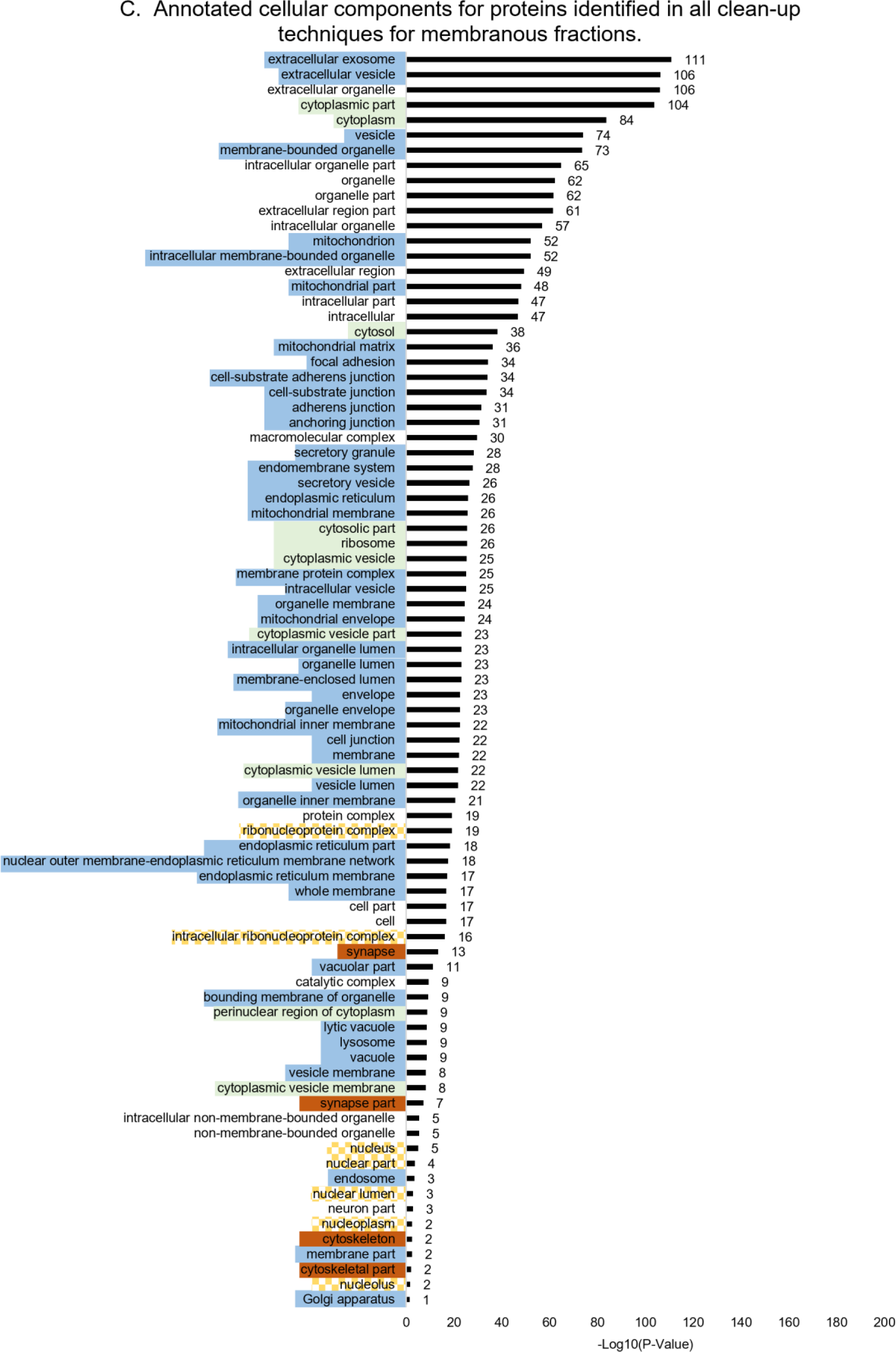
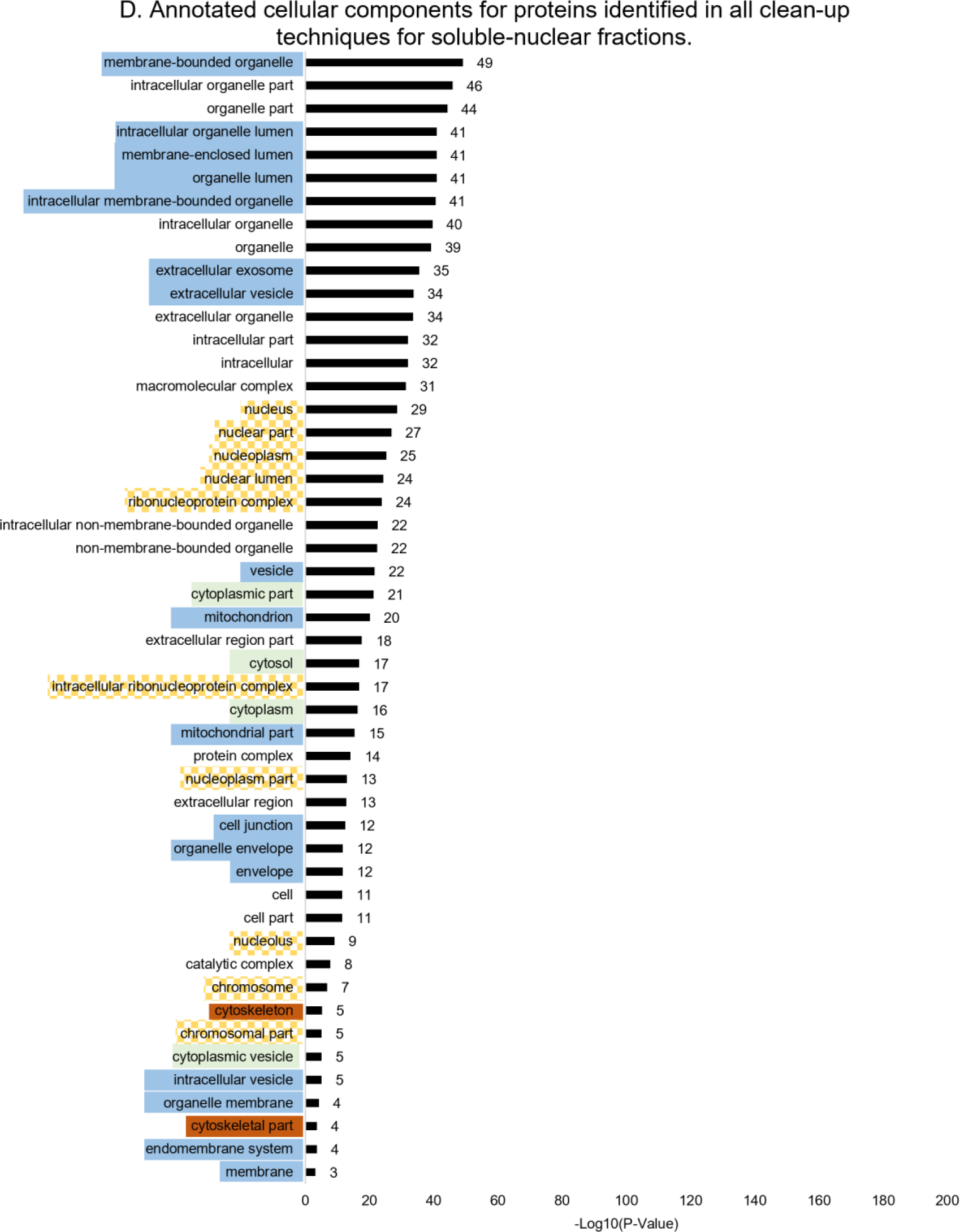
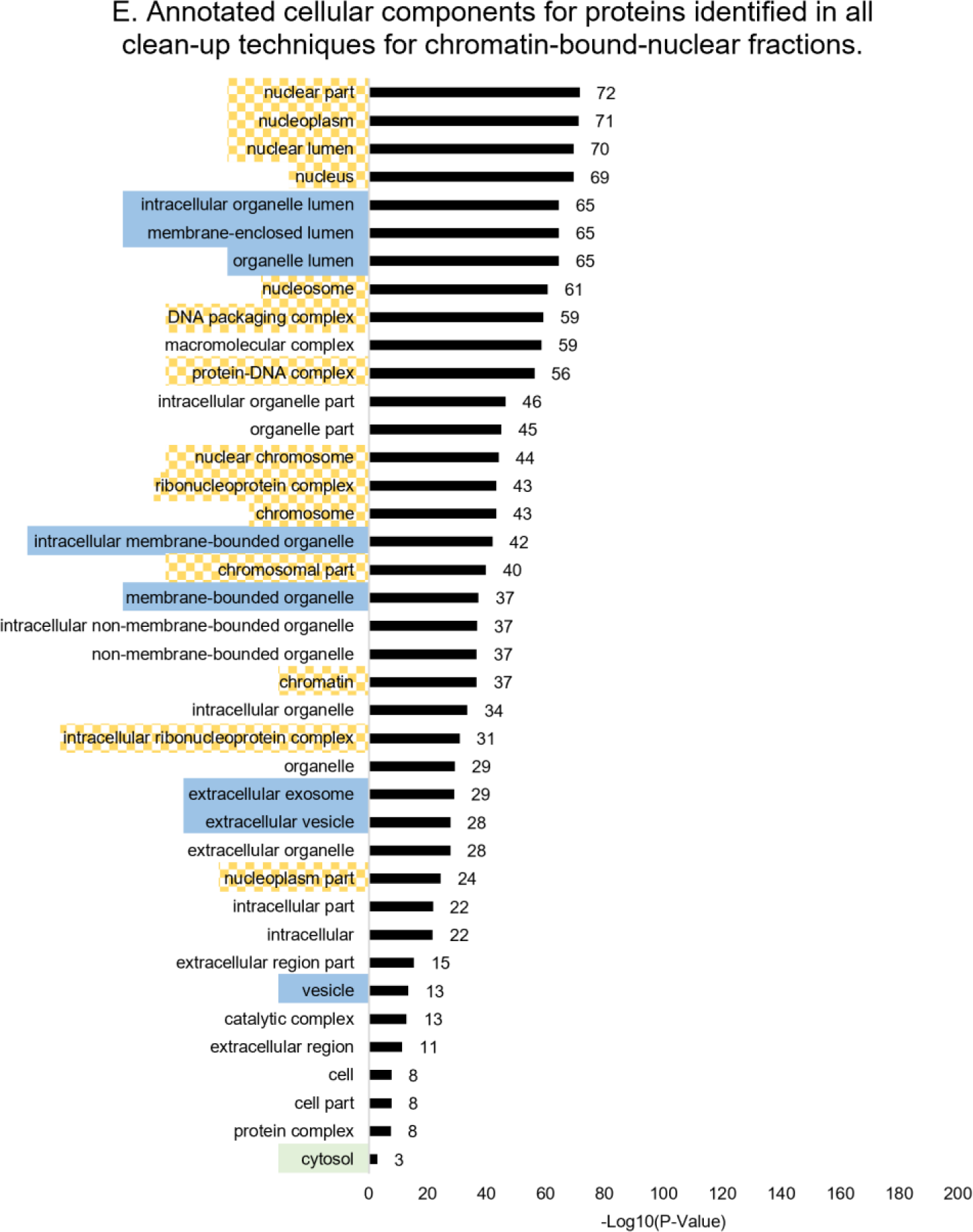
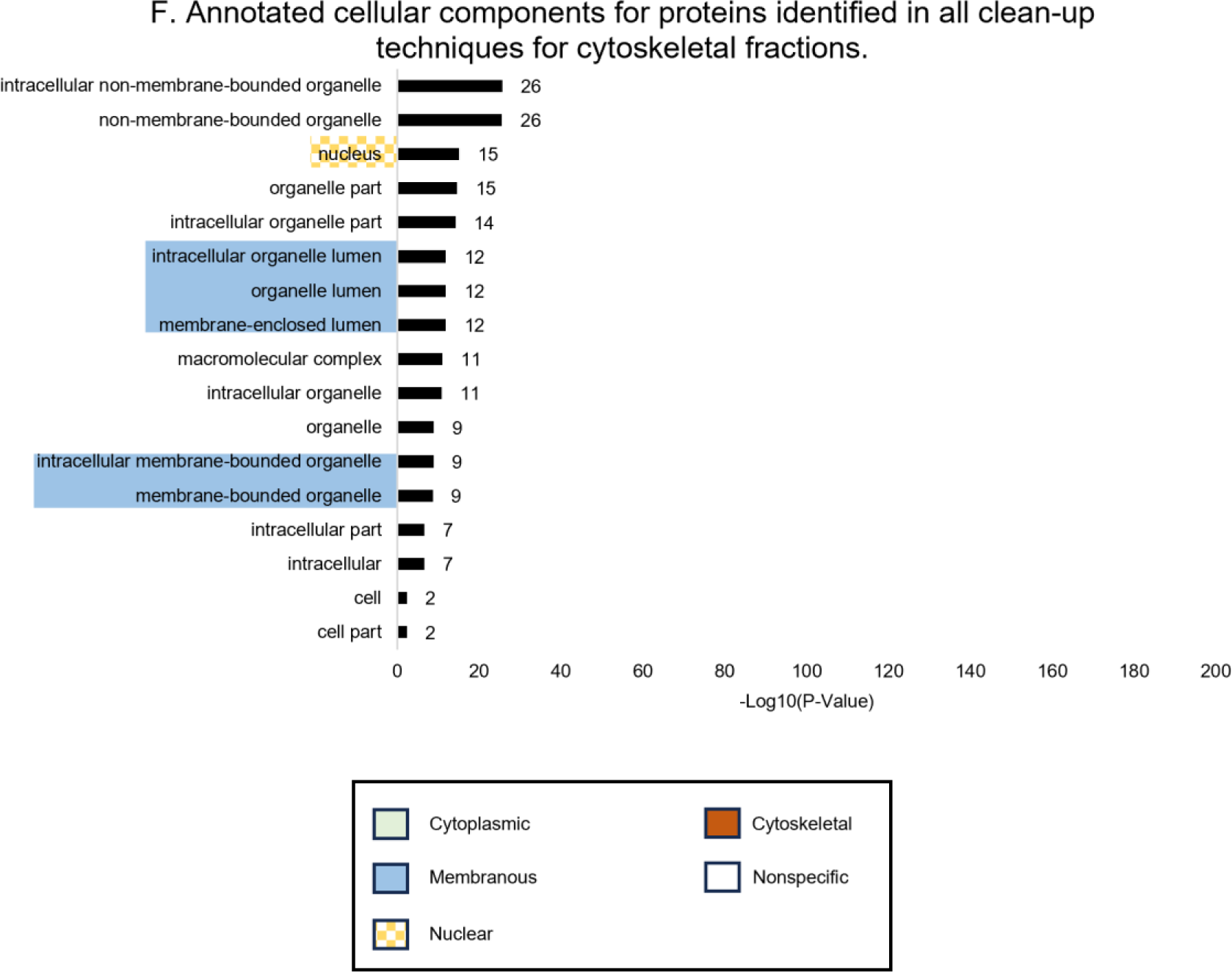
Annotated cellular components of proteins identified by all clean-up techniques. Cellular components output from DAVID functional annotation tool for proteins identified in all clean-up techniques in whole-cell lysates (A), cytoplasmic (B), membranous (C), soluble-nuclear (D), chromatin-bound-nuclear (E), and cytoskeletal (F) fractions. Cellular components are grouped into cytoplasmic (green highlighted), membranous (blue highlighted), nuclear (yellow-checkered highlighted), cytoskeletal (red highlighted), or nonspecific (not highlighted).

### Figures S8A-F Methods

The Database for Annotation, Visualization, and Integrated Discovery (DAVID) functional annotation tool^30^ was used to annotate proteins to cellular components. Uniprot accession numbers for proteins identified by all four clean-up methods (Figure 2) were input into DAVID as a gene list. Homo sapiens was selected as the species, and the functional annotation tool was used to output a functional annotation chart for GOTERM_CC_ALL. The count was set to 50 and ease to 0.05 with a Benjamini correction applied to the options tab. Cellular-component outputs from DAVID were manually grouped into cytoplasmic, membranous, nuclear, cytoskeletal, or nonspecific categories using Uniprot subcellular localization,^5^ Alliance of Genome Resources (v.5.4.0),^6^ and the Gene Ontology Resource (Release 2023-06-11).^7,8^ Nonspecific-cellular components include ‘cell’, ‘organelle’, or any cellular component that could not fit in one of the other categories.

**Figures S9A-F.**
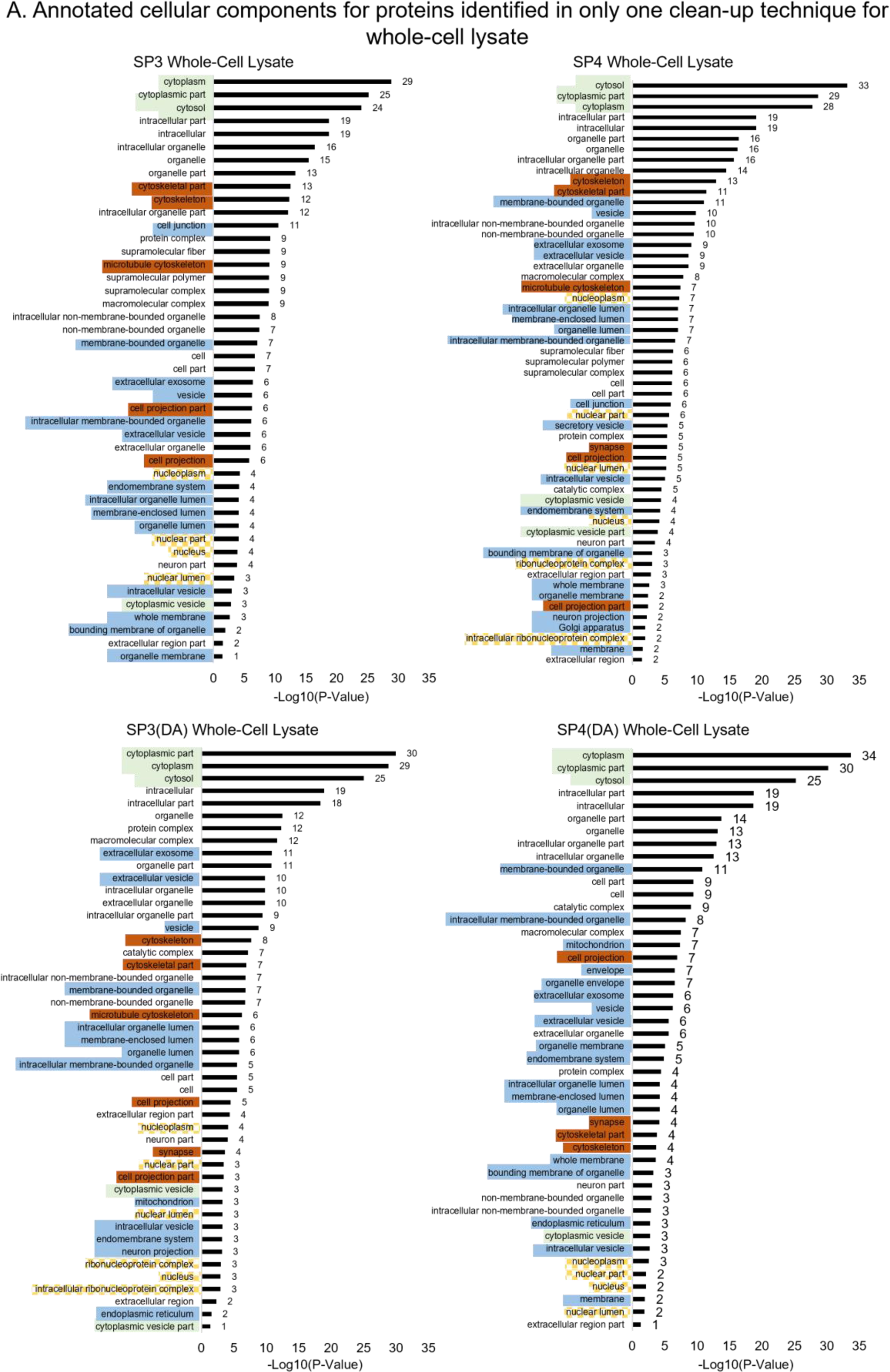
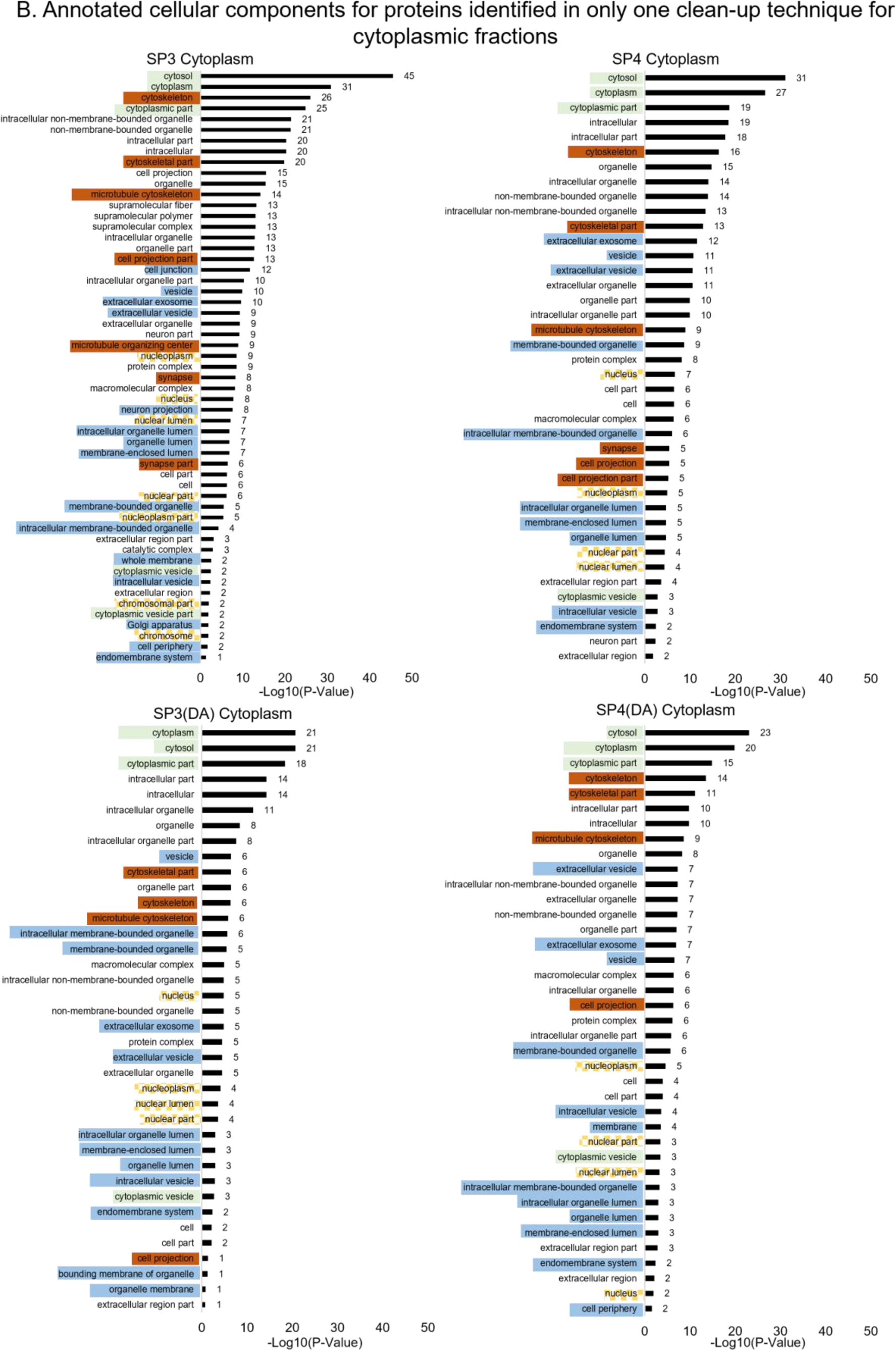
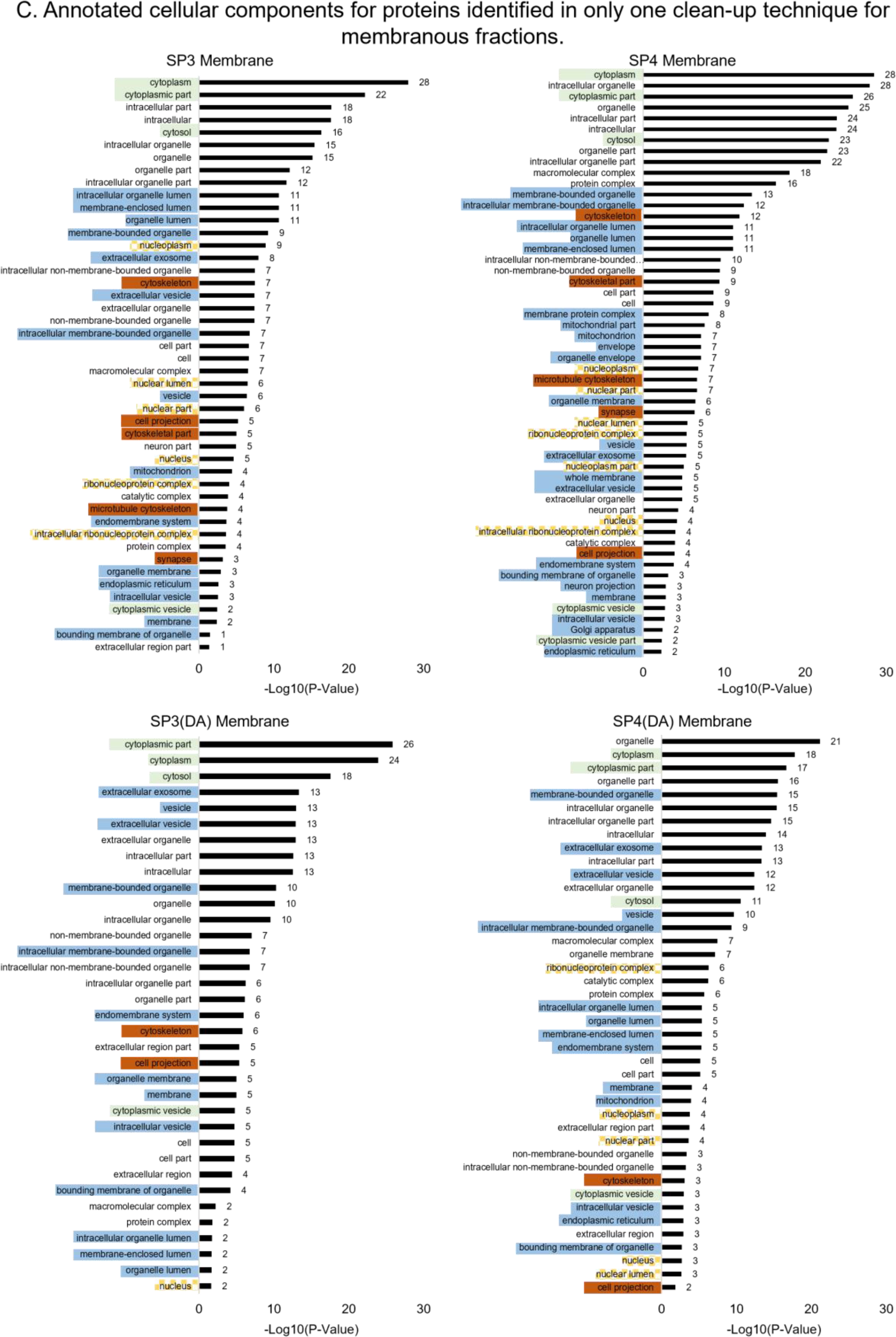
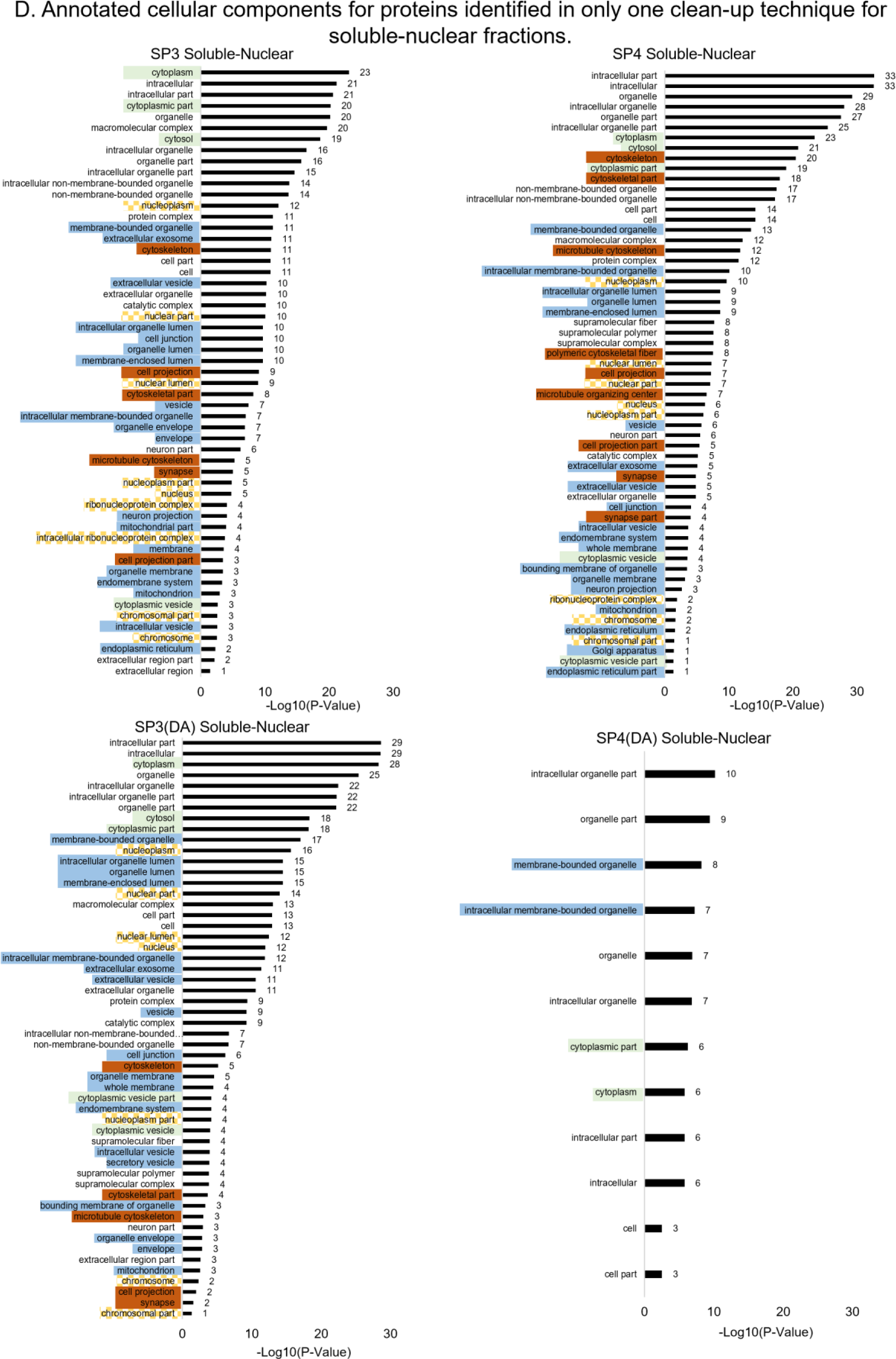
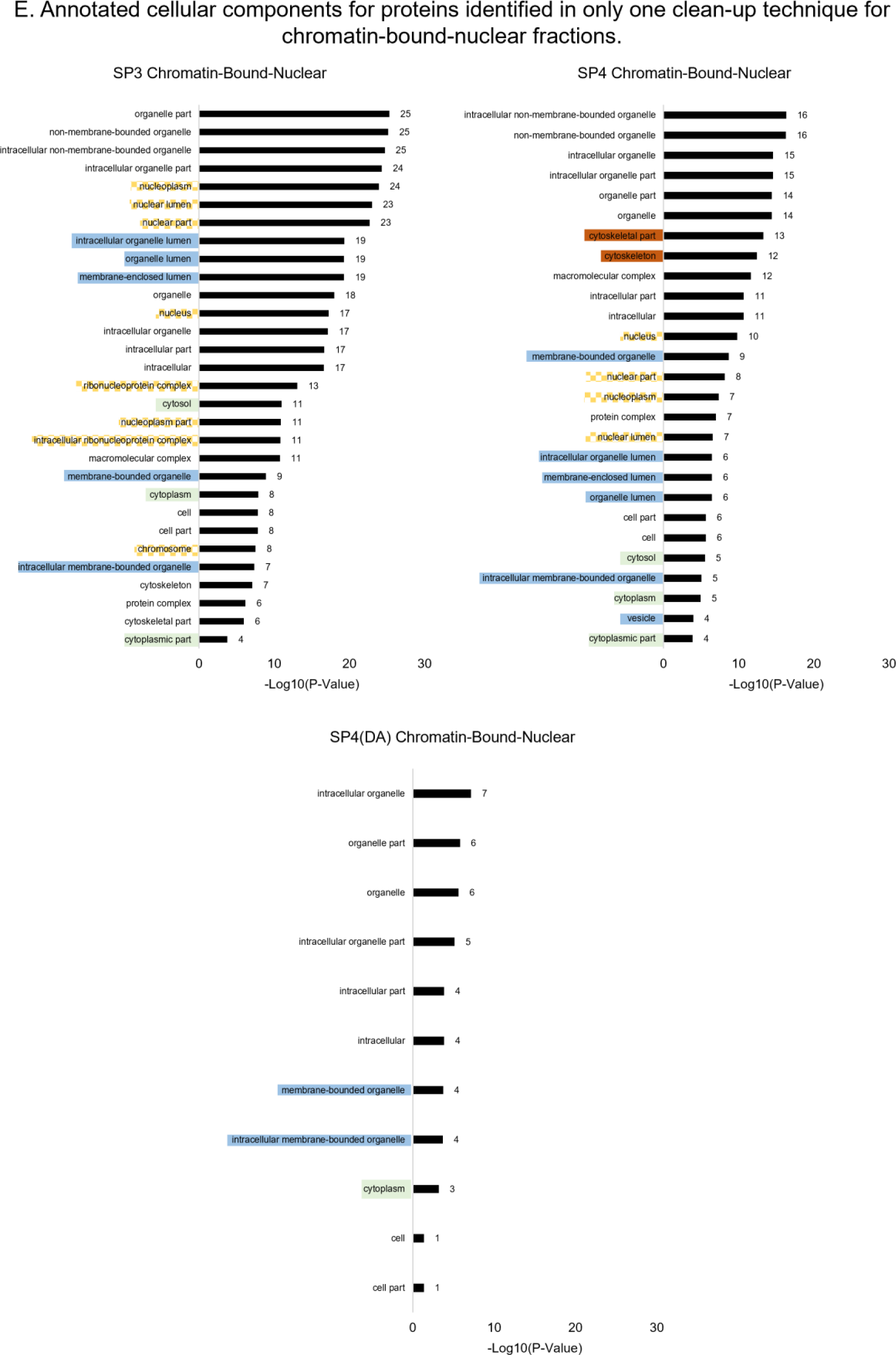
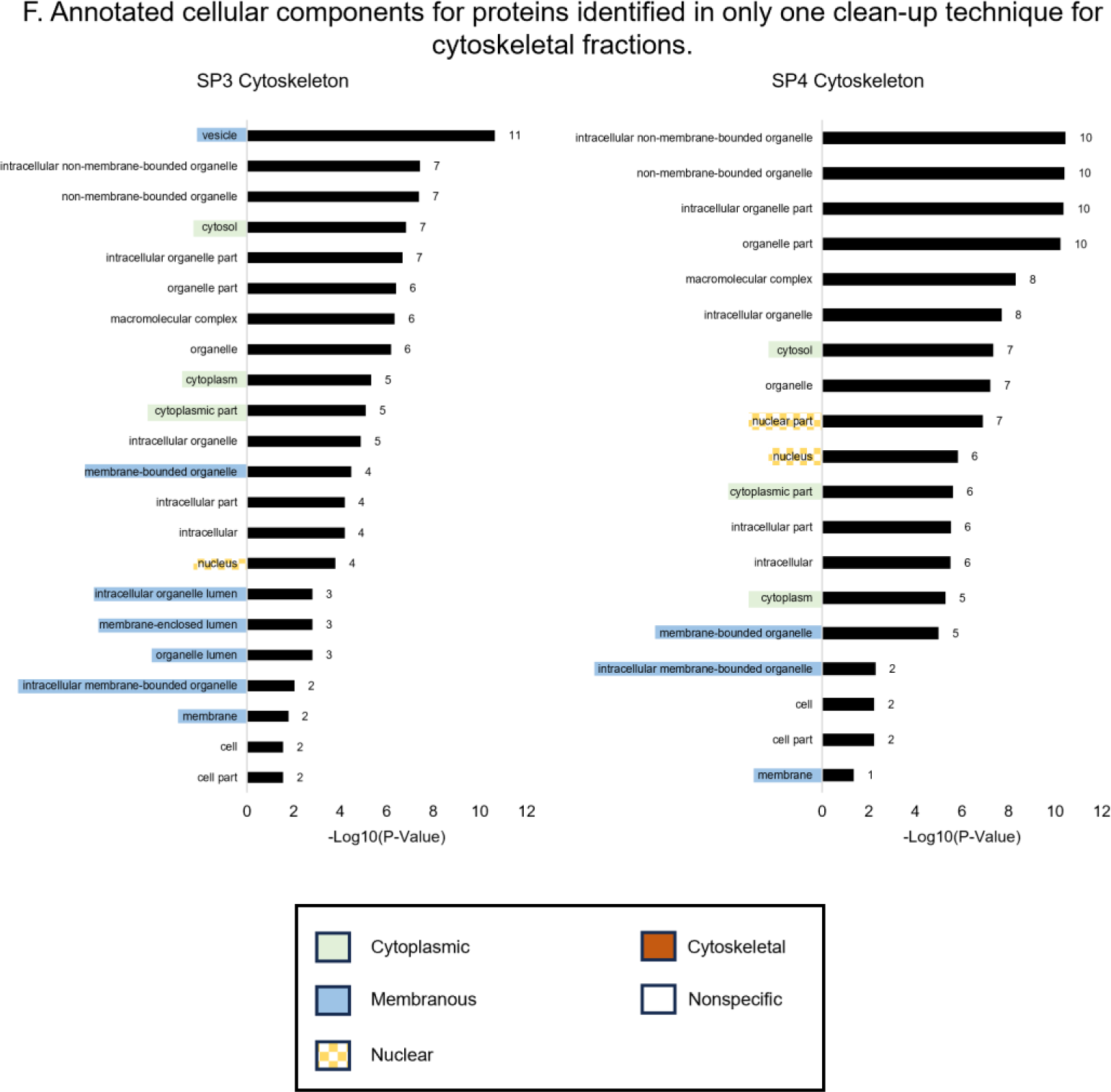
Annotated cellular components of proteins identified using only one clean-up technique. Cellular components output from DAVID functional annotation tool for proteins identified in only one clean-up technique for whole-cell lysates (A), cytoplasmic (B), membranous (C), soluble-nuclear (D), chromatin-bound-nuclear (E), and cytoskeletal (F) fractions. Cellular components are grouped into cytoplasmic (green highlighted), membranous (blue highlighted), nuclear (yellow-checkered highlighted), cytoskeletal (red highlighted), or nonspecific (not highlighted).

### Figures S9A-F Methods

The Database for Annotation, Visualization, and Integrated Discovery (DAVID) functional annotation tool^30^ was used to annotate proteins to cellular components. Uniprot accession numbers for proteins identified in only one clean-up technique (Figure 2) were input into DAVID as a gene list. Homo sapiens was selected as the species, and the functional annotation tool was used to output functional annotation chart for GOTERM_CC_ALL. The count was set to 50 and ease to 0.05 with a Benjamini correction applied to the options tab. Cellular components output from DAVID were manually grouped into cytoplasmic, membranous, nuclear, cytoskeletal, or nonspecific categories using Uniprot subcellular localization,^5^ Alliance of Genome Resources (v.5.4.0),^6^ and the Gene Ontology Resource (Release 2023-06-11 http://geneontology.org/).^7, 8^ Nonspecific-cellular components include ‘cell’, ‘organelle’, or any cellular component that could not fit in one of the other categories.

**Figure S10.**
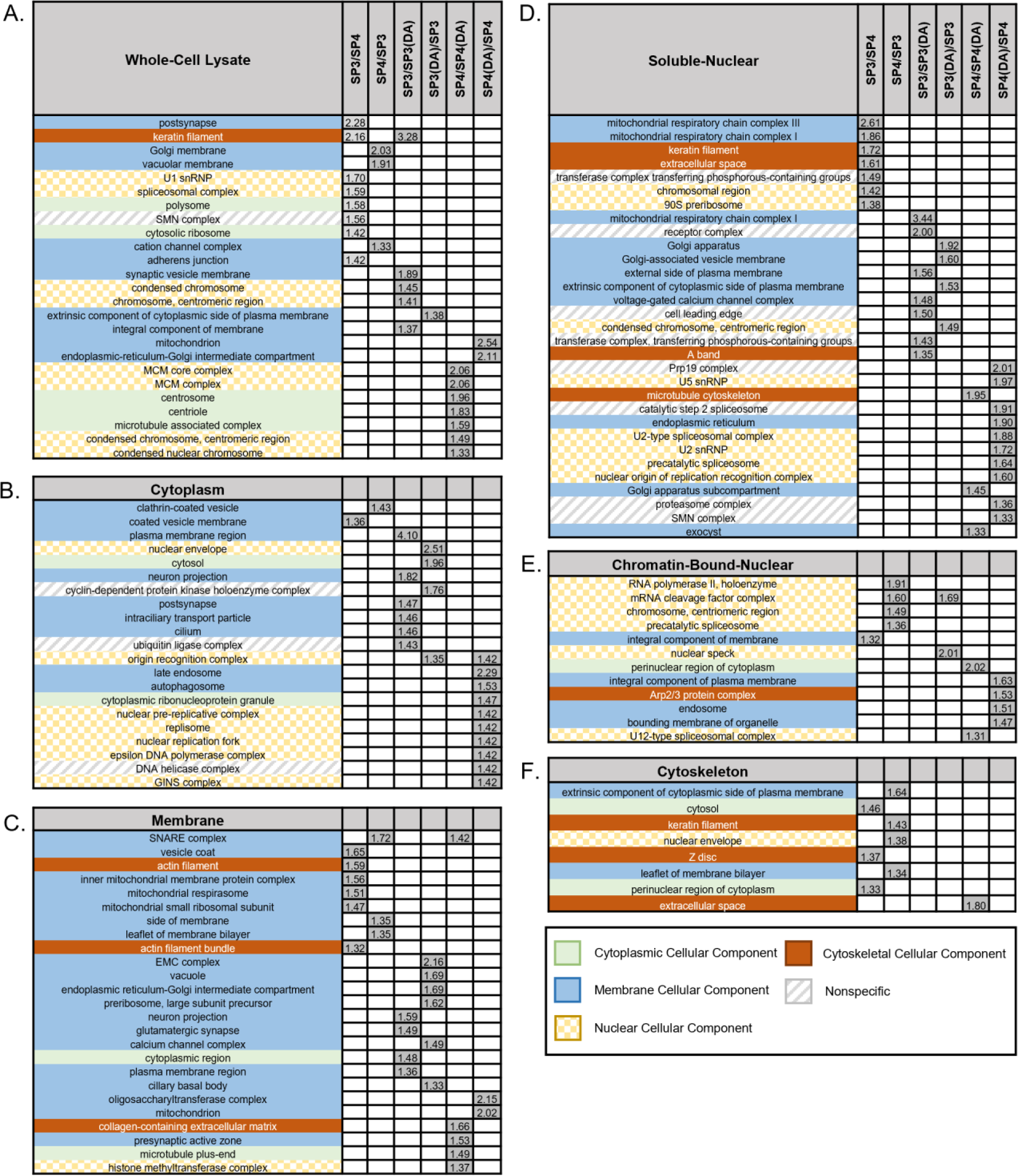
Gene ontology-Slim analysis for all samples. Comparison of gene ontology-Slim (GO-Slim) cellular components for enriched proteins from whole-cell lysate (A), cytoplasmic (B), membranous (C), soluble-nuclear (D), chromatin-bound-nuclear (E), and cytoskeletal (F) fractions. Enriched proteins were identified by volcano plots (Figure 4). Columns show comparisons of enriched cellular components for SP3 versus SP4 (SP3/SP4, i), SP4 versus SP3 (SP4/SP3, ii), SP3 versus SP3(DA) (SP3/SP3(DA), iii), SP3(DA) versus SP3 (SP3(DA)/SP3, iv), SP4 versus SP4(DA) (SP4/SP4(DA), v), and SP4(DA) versus SP4 (SP4(DA)/SP4, vi). White-colored boxes indicate that a cellular component ontology was not significantly enriched by either method in the comparison. Grey-colored boxes represent -log10(P-values) for cellular components that were significantly enriched in one clean-up method versus the other via GO-Slim analysis (P-value < 0.05 in PANTHER).

### Figure S10 Methods

Pair-wise comparisons were searched in Progenesis QIP and proteins were selected to be “exported to pathways tool” on the “Review Proteins” tab in Progenesis, and the PANTHER classification system was selected as the pathways analysis tool. The statistical enrichment test was used to compare pairwise comparisons (SP3 versus SP4, SP3 versus SP3(DA), and SP4 versus SP4(DA)). The proteins were exported via a .txt file that was imported into PANTHER (Version 17.0, accessed February 2023). Homo sapiens were selected as the organism and a statistical enrichment test was selected with “PANTHER GO-Slim Cellular Component”. Gene ontologies were determined in PANTHER by assessing the genes associated with enriched proteins by each clean-up technique. Gene products are then correlated to cellular-component ontologies (in PANTHER) by determining the location that the gene occupies when it carries out a molecular function.^9^ Enriched cellular components with P-values < 0.05 were used to construct tables in Figures 6 and S10.

### Supplementary Tables

**Table S1.**
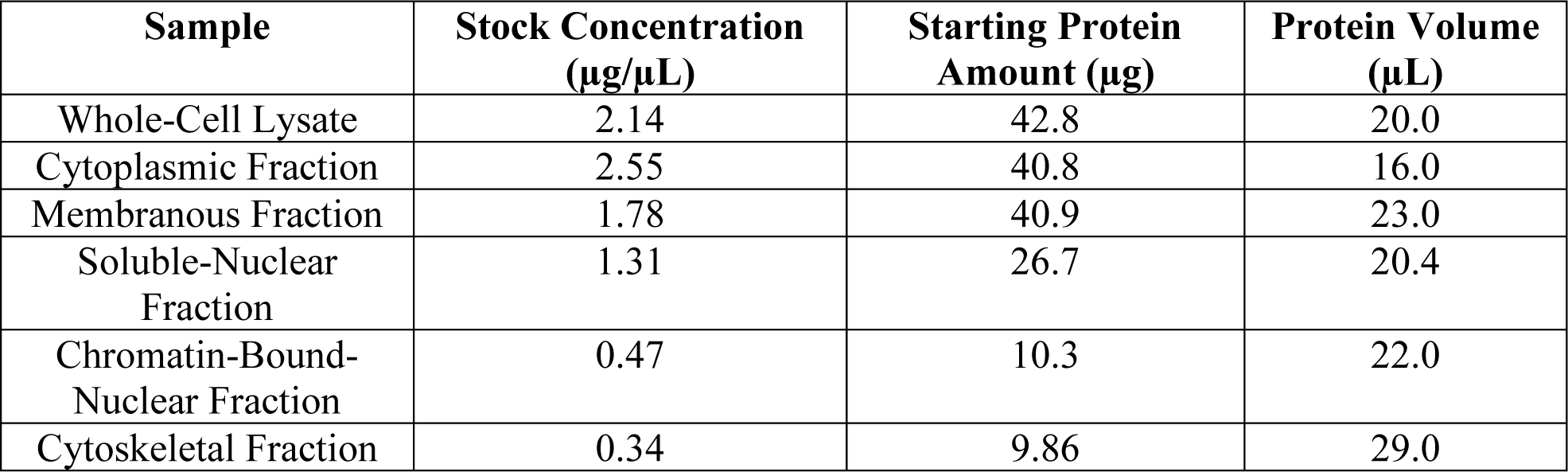
Starting protein amounts and concentrations.

| **Sample** | **Stock Concentration**<br>**($\mu\text{g}/\mu\text{L}$)** | **Starting Protein**<br>**Amount ($\mu\text{g}$)** | **Protein Volume**<br>**($\mu\text{L}$)** |
| --- | --- | --- | --- |
| Whole-Cell Lysate | 2.14 | 42.8 | 20.0 |
| Cytoplasmic Fraction | 2.55 | 40.8 | 16.0 |
| Membranous Fraction | 1.78 | 40.9 | 23.0 |
| Soluble-Nuclear<br>Fraction | 1.31 | 26.7 | 20.4 |
| Chromatin-Bound-Nuclear Fraction | 0.47 | 10.3 | 22.0 |
| Cytoskeletal Fraction | 0.34 | 9.86 | 29.0 |

Protein concentrations in stock solutions of whole-cell lysates and subcellular fractions were determined using bicinchoninic acid (BCA) assays (ThermoFisher Scientific, Waltham, MA). Aliquots were stored at -80 °C until sample preparation and LC-MS/MS analysis. SP3 has previously been shown to be an ideal clean-up technique for protein samples prepared at concentrations lower than 0.25 µg/µL.^1^ Thus, the starting protein concentrations for these fractions were all greater than 0.25 µg/µL to ensure that one method was not biased over another based on the starting protein concentration.

**Table S2.**
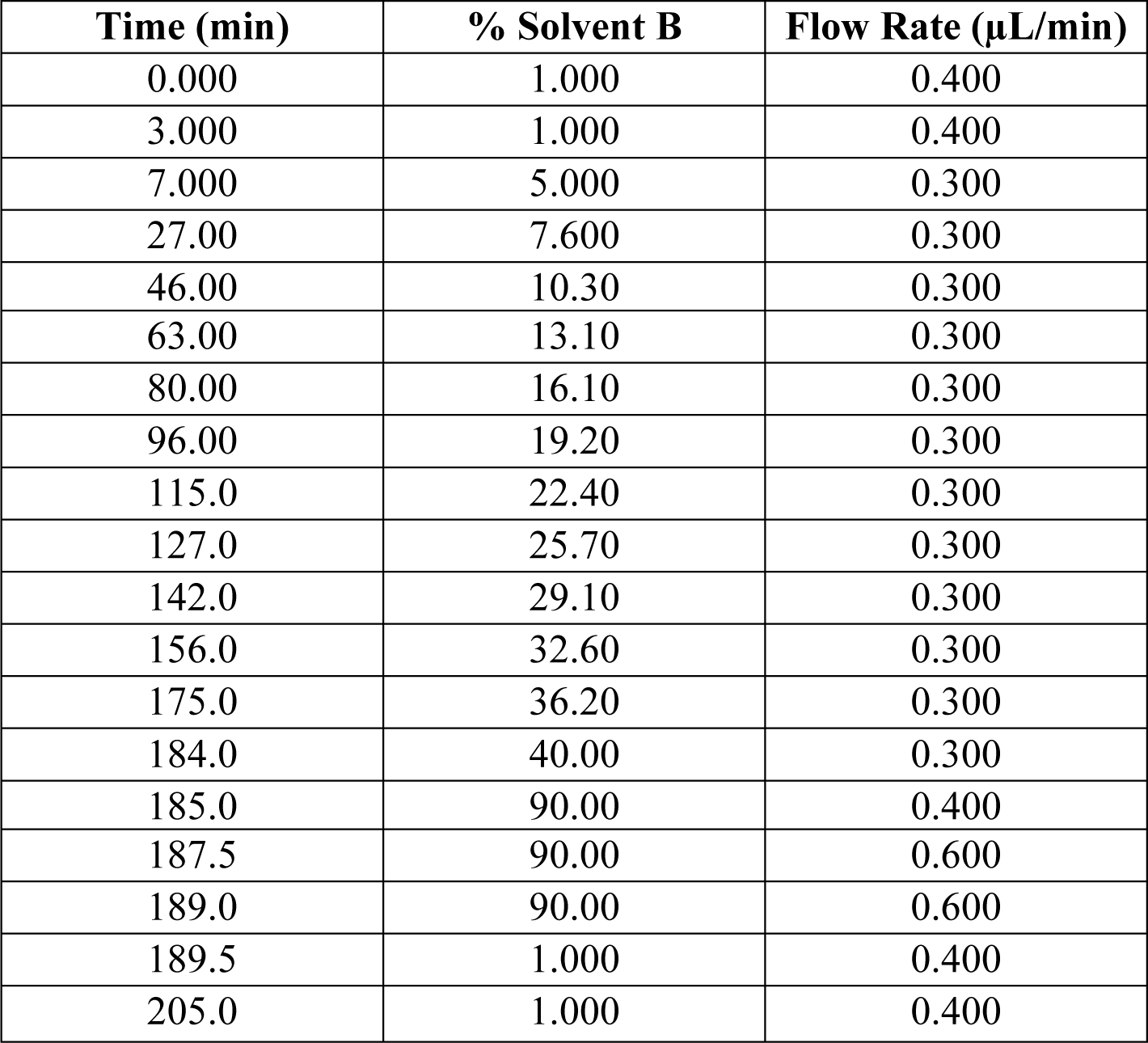
Liquid Chromatography Gradient.

A gradient composed of solvent A (99.9% water and 0.1% formic acid) and solvent B (99.9% acetonitrile and 0.1% formic acid) was used to separate peptides prior to MS analysis. Gradient was adapted from a previous publication.^3^

**Table S3.** Optimized Synapt G2-S HDMS Tune Page Parameters. Tune parameters were optimized from previous literature^29^ to enable mobility separation in the Triwave cell.

| **MS Tune Page Parameter** | **Value** |
| --- | --- |
| **Nanoflow** |  |
| Capillary Voltage | 3.2 kV |
| Sampling Cone | 40 V |
| Source Offset | 80 V |
| Source Temperature | 100 °C |
| **Step Wave** |  |
| Step Wave 2 |  |
| Wave Velocity | 20 m/s |
| Wave Height | 15 V |
| Source Ion Guide |  |
| Wave Velocity | 2700 m/s |
| Wave Height | 5 V |
| **Trapping** |  |
| Mobility Trapping |  |
| Release Time | 500 μs |
| Trap Height | 15 V |
| Extract Height | 0 V |
| Mobility Delay |  |
| Additional Bins | 3 |
| IMS Wave Delay | 450 μs |
| **TriWave** |  |
| IMS Wave Velocity | 850 m/s |
| IMS Wave Height | 40 V |
| Transfer Wave Velocity | 190 m/s |
| Transfer Wave Height | 5 V |
| **IMS-Config** |  |
| Start Wave Velocity | 1200 m/s |
| End Wave Velocity | 850 m/s |
| Ramp Over Full IMS Cycle | ON |

